# *BCL10* Gain-of-Function Impairs Germinal Center Participation by Naïve B Cells and Cooperates with *BCL6* to Drive Extrafollicular Diffuse Large B-Cell Lymphoma

**DOI:** 10.64898/2026.11.03.764037

**Authors:** Dhanvantri Chahar, Abdessamad A. Youssfi, Caroline A. Coughlin, Hsuan-Ting Huang, Vivian L.S. Kuan, Venu Venkatarame Gowda Saralamma, Marianna Lekakis, Dylan Huynh, Francesco Maura, Catalina Amador, Daniel Bilbao, Jayanta Chaudhuri, Jonathan H. Schatz

## Abstract

*BCL10* gain-of-function (GoF) mutations occur in diffuse large B-cell lymphoma (DLBCL) cases of the BN2/C1 molecular subgroup. These alterations activate CARD11-BCL10-MALT1 (CBM) signaling and drive resistance to multiple drug classes, but their role in disease pathogenesis is incompletely understood. In particular, interactions between BCL10 GoF and additional DLBCL oncogenes are undefined. Here, we show BCL10 GoF in naïve B cells blocks their participation in germinal center (GC) reactions and permits transformation to DLBCL with a non-GC B-cell phenotype. Most clinical cases with BCL10 GoF also carry translocation of the *BCL6* proto-oncogene, and we show that co-activation of these two lesions together promotes pronounced oncogenic synergy. Detailed single-cell analyses of underlying premalignant B-cell populations revealed expansion of extrafollicular memory B cells whose gene-expression program remains prominent in resulting lymphomas in both the mouse and corresponding human tumors. We therefore define a novel role for *BCL10* activating mutations in enabling *BCL6* lymphomagenesis.

**Statement of Significance:** Complex mutational patterns underlie molecular subtypes of DLBCL, the most common lymphoma. We define the basis for synergy between co-deregulated oncogenes in a disease cluster commonly arising from non-GC B-cells. Extrafollicular memory B cells are implicated as the precursors undergoing malignant transformation by the dual deregulation of *BCL6* and *BCL10*.

## Introduction

*BCL10* gain-of-function (GoF) mutations occur in diffuse large B-cell lymphoma (DLBCL) cases with *BCL6* translocations and/or *NOTCH2* mutations, known as BN2 by LymphGen or Cluster 1 by DLB*class* (BN2/C1) (1–4). These mutations activate CARD11-BCL10-MALT1 (CBM) signaling independently of upstream BTK-mediated B-cell receptor (BCR) engagement and promote decreased responses to multiple classes of targeted drugs (5,6). We recently showed that lymphoma cells engineered with these alterations activate protection from apoptosis downstream of constitutive NF-kB pathway activation (5). Accordingly, BCL2 inhibition with the clinical compound venetoclax restored sensitivity of these systems to BTK inhibition. Accurate in vivo models of BCL10 mutational lymphomagenesis, however, are lacking. Moreover, their roles in pathogenesis, in particular their relationship to defining BN2/C1 molecular alterations, remain unclear. Understanding these dynamics is crucial for developing effective therapies.

BCL6 is a master regulator of the germinal center (GC) reaction in antigen-responsive B cells, functioning as a transcriptional repressor that promotes and maintains the reaction (7,8). In the GC dark-zone (DZ), cells proliferate rapidly and acquire mutations to the immunoglobulin heavy chain variable region (IgHV), which are subsequently tested for impact on antibody binding affinity to antigens in the light zone (LZ) (9,10). By selecting for advantageous IgHV mutations and eliminating others, GCs result in the generation and propagation of both plasma and memory B-cell populations expressing mature antibodies with optimized affinity against specific antigens (11). When perturbed, however, GCs also underly the majority of mature B cell-derived lymphomas, including DLBCL (8). *BCL6* in particular is a proto-oncogene frequently over-expressed in DLBCL through the translocations that co-define BN2/C1 cases and a variety of other mechanisms (12,13). BCL6 promotes rapid proliferation, protection from apoptosis, and tolerance of DNA damage (14), among other potentially oncogenic functions, but its activation alone is generally inadequate for lymphomagenesis (15). *Iµ-HABCL6* mice, which recapitulate the human t(3;14)(q27;q32) *BCL6* translocation, develop late-onset lymphomas with incomplete penetrance, consistent with a requirement for cooperating drivers (16). Few studies exist evaluating cooperation in lymphomagenesis between BCL6 deregulation and other recurrently modified genes that cluster in BN2/C1 cases.

Here, we generated conditional transgenic mice that express an activating BCL10 truncation (S136X) upon co-expression of cre recombinase. *BCL10*^S136X^ expression in naïve B (NB) cells drove co-stimulation-independent proliferation in response to BCR engagement and blocked participation in GC reactions amid inflamed microenvironments with prominence of exhausted T cells. Non-GCB DLBCL arose in secondary recipients after isolation of B-cell clones with malignant potential, but crossing these animals with *Iµ-HABCL6* promoted the disease in 100% of primary animals. Single-cell investigation of pre-malignant B-cell populations revealed expansion of extrafollicular IL9R-expressing (M3) memory B cells (MBC) driven by the co-expression of *BCL10*^S136X^ and *BCL6*. Correspondingly, the M3 MBC gene-expression program was prominent in clinical cases with co-occurrence of *BCL10* GoF mutation and *BCL6* translocation. We therefore demonstrate oncogenic synergy between *BCL10* mutational activation and *BCL6* deregulation in DLBCL, arising from extrafollicular MBCs, and reflecting observations in many clinical cases.

## Results

### Expression of an activating BCL10 truncation in NB cells facilitates transformation to non-GCB-DLBCL

To assess BCL10 GoF in vivo, we engineered the activating S136X truncation (5,6) with GFP co-expression into the murine *Rosa26* locus with a preceding Lox-STOP-Lox cassette **(Fig. 1A).** These *Rosa26*-LSL-*BCL10*^trunc^ (RSB) animals are born at expected numbers and ratios by sex and have normal lifespans and phenotype. Cre introduction to murine embryonal fibroblasts (MEFs) induced expression of truncated BCL10 protein and eGFP **(Supplementary Fig. 1A).** We bred RSB^+/−^ to Mb1-Cre^+/−^ animals (17) and confirmed recombination of the transgenic allele occurs only in double-heterozygous *Mb1.BCL10*^trunc^ offspring (**Supplementary Fig. 1B-C)**. These animals had GFP expression in circulating B cells not seen in littermate genotypes and truncated BCL10 protein detectable in purified splenic B cells (**Supplementary Fig. 1D-E**). RSB therefore permitted lineage-specific *BCL10*^trunc^ induction, with no evidence of expression in an absence of Cre. Bone marrow showed induction of transgene expression beginning at the early pro-B stage as expected under the Mb1 promoter (17) (**Fig. 1B**). Expression of a *CARD11* activating mutation at the same point of B-cell development promoted rapid onset of aggressive lymphomas (18). When we followed *Mb1.BCL10*^trunc^ animals compared to littermates, however, we found no survival differences in two years (**Fig. 1C**). Phenotyping at necroscopy revealed no clear evidence of lymphoma or leukemia, and serial flow cytometry of peripheral blood showed no consistent increase in circulating B cells (**Supplementary Fig. 1F**). Oncogenesis from transgenic B cells may emerge in secondary recipients after passage ex vivo or through immunodeficient (e.g. *Rag*-deficient) animals to select clones with malignant potential (19–21). We therefore cultured cells from lymph nodes of primary animals on irradiated murine fibroblast feeder layers (**Fig. 1D**). A GFP^+^ B-cell population emerged during serial passage of *Mb1.BCL10*^trunc^ cells, while littermate genotypes produced no populations that proliferated ex vivo. We introduced 2.5 x 10^5^ of these cells by tail-vein injection to irradiated (3.5 Gy) WT B6 recipients, which become moribund with palpable splenomegaly ∼12 weeks after injection (**Fig. 1E**). Pathologic evaluation revealed GFP^+^ large B-cell lymphomas involving spleen and lymph nodes. Tumors were CD10-negative IRF4/MUM-1^+^ LBCLs that stained strongly for BCL6, recapitulating non-GCB clinical cases by the Hans algorithm (**Fig. 1F, Supplementary Fig. 1G**) (22) and demonstrating B-cell monoclonality (**Supplementary Fig. 1H**). These tumors were readily transplanted to further recipients, demonstrating cell-dose dependent tumor onset (**Fig. 1G**). Disease phenotypes were the same, including splenomegaly (**Fig. 1H**) and histologic features (not shown). FACS analysis of recipient splenocytes showed predominance of enlarged B lymphocytes with bright expression of CD19 and IgM, loss of B220 and IgD, and no evidence of plasma-cell differentiation (CD138^−^, **Fig. 1I-K**). *BCL10*^trunc^ expression in pre-GC B cells therefore facilitated transformation to DLBCL with non-GCB features.

**Figure 1:**
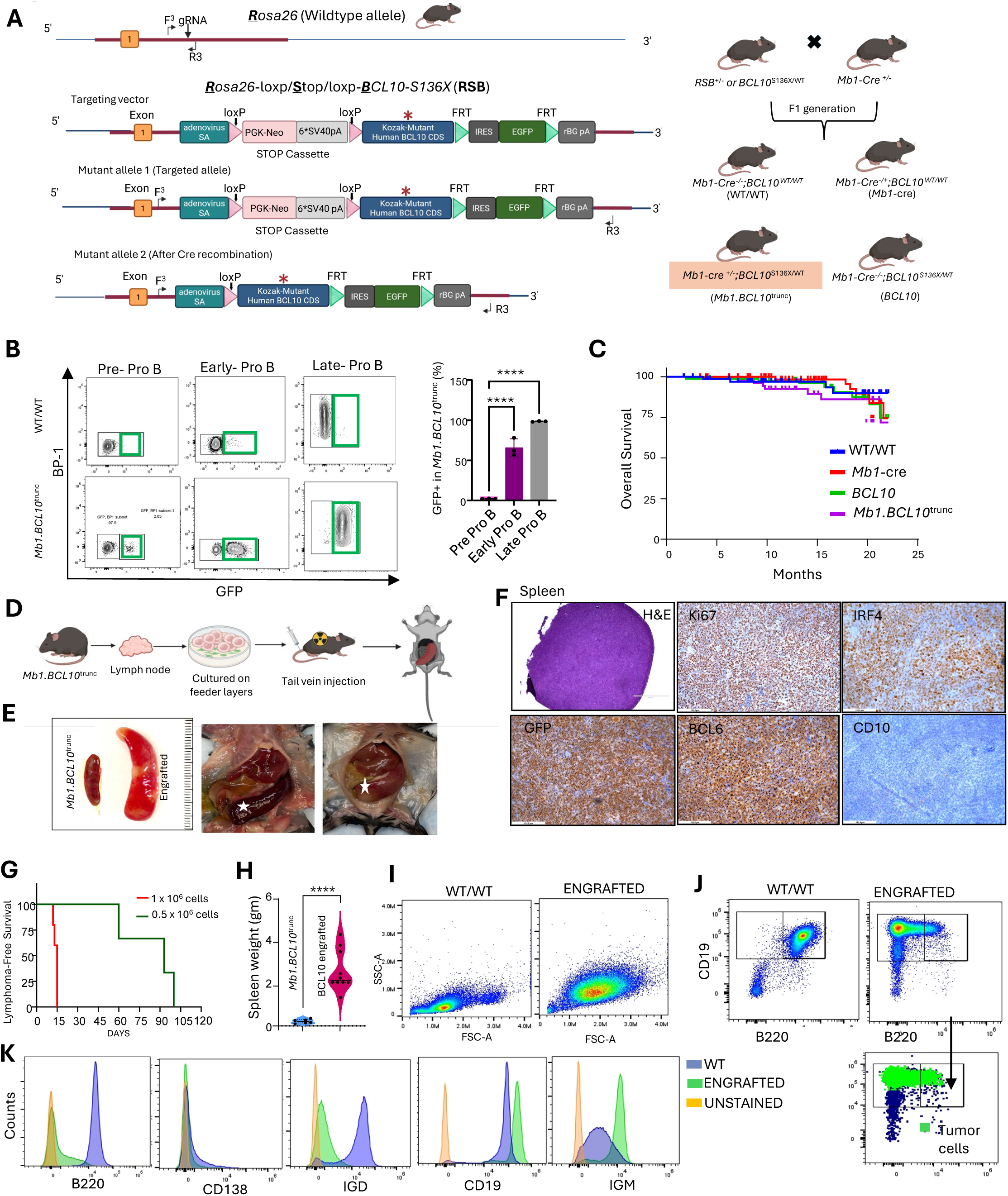
Cre-inducible expression of activating BCL10 truncation in NB cells. **A**. (Left) Targeting strategy at *ROSA26* locus. A loxP-STOP-loxP-*BCL10*-S136X-IRES-EGFP cassette (RSB) (red asterisks indicate S136X) was knocked into Rosa26 locus. The STOP cassette (PGK-Neo,6xSV40pA) blocks transcription until Cre-mediated excision, after which mutant *BCL10* and EGFP are expressed from the endogenous *Rosa26* promoter. F3 and R3 are genotyping primers; FRT site flank IRES-EGFP. (Right) Breeding scheme showing RSB+/−were bred to Mb1-cre+/− to restrict expression to B cells, generating Mb1-cre^+/−^;BCL10^S136X/WT^ (shaded) and three littermate control genotypes. (Mb1-cre^+/−^;BCL10^S136X/WT^ was annotated as *Mb1.BCL10*^trunc^ throughout the text and figures in the paper). **B**. (Right) Representative flow plots showing proper recombination of mutant allele and expression of GFP in Pro-B cells in bone marrow population of *Mb1.BCL10*^trunc^ mice compared to WT/WT (n=3). Green gates mark the EGFP^+^ population. EGFP serves as a marker for Cre-mediated excision of the STOP cassette and thus for mutant *BCL10*-S136X expression. (Left) Quantification of the EGFP^+^ population. **C**. Overall survival of unimmunized cohort comparing *Mb1.BCL10*^trunc^ to littermate controls. **D**. Schematic diagram of transfer strategy for lymphoma cells derived from *Mb1.BCL10*^trunc^. **E**. Representative images of spleen from normal mice (WT/WT) and the mice with transplanted B cells derived from *Mb1.BCL10*^trunc^ lymph nodes. Gross dissection demonstrates the huge splenomegaly (marked with white asterisk) in multiple mice post engraftment. **F**. H&E and IHC images from engrafted mice spleen; Ki67, GFP (correlates human *BCL10*^S136X^), CD10, IRF4 and BCL6. **G**. Survival curve demonstrates an inverse correlation between the number of transplanted tumor cells and mouse survival. **H**. Quantification of spleen weights (gm) of age matched control *Mb1.BCL10*^trunc^ mice (blue) and recipient WT mice engrafted with *Mb1.BCL10*^trunc^ derived lymphoma cells measured at endpoint post engraftment. Engrafted mice developed marked splenomegaly relative to controls. (n=10 mice/engrafted group). ****p<0.0001. **I**. Altered cell size and granularity in engrafted mice spleen cells. Representative forward scatter (FSC-A) versus side scatter (SSC-A) density plots of splenocytes from WT/WT control (left) and engrafted mice (right). Cells from engrafted animals showed shift toward higher FSC-A and SSC-A showing the shift in the scattering plot of tumor B cells indicative of large B cells typical of transformed lymphoblasts. **J**. Expansion of a B220 ^low/int^, CD19^+^ population in engrafted mice. Representative CD19 versus B220 plots showing a higher shift in the CD19 expression of the tumor B cells (green) isolated from spleen of engrafted mice compared to endogenous B cells (blue) within the engrafted mice. GFP (correlates human *BCL10-*S136X) shows the tumor cells in the B cell population. **K**. Surface marker phenotype of engrafted lymphoma cells. Representative histograms overlays of B220, CD138, IgD, CD19and IgM expression on cells from WT/WT control mice (blue) and engrafted mice (green) with unstained controls (orange) shown for each marker. Cells were gated on viable cells.

### BCL6 Synergizes with BCL10 GoF in Lymphomagenesis

Given high BCL6 expression in *BCL10*^trunc^ tumors, we hypothesized BCL6 and BCL10 GoF cooperate in lymphomagenesis. Examining clinical cases revealed that *BCL6* rearrangements and *BCL10* activating mutations frequently co-occur, impacting 15.7% of DLB*class* C1 and 28.3% of LymphGen BN2 cases, associated with increased frequency of unclassified COO in both (**Fig. 2A-B**). Cases with both lesions (n=18) in DLB*class* had similar overall survival compared to the rest of the cohort and to other C1 cases (**Fig. 2C, Supplementary Fig. 2A**) (2,4). To model co-occurrence, we crossed *Mb1.BCL10*^trunc^ animals with *Iµ-HABCL6* (16). *Iµ-HABCL6* activates BCL6 expression in mature B cells before GC entry, allowing us to test cooperation with *BCL10*^S136X^ pre- and post-GC activation (23). *BCL6.Mb1.BCL10*^trunc^ triple heterozygous animals developed onset of fatal lymphomas with 100% penetrance between ages 3 and 20 months (**Fig. 2D**), contrasting with late onset and incomplete penetrance in *Iµ-HABCL6*, consistent with previous observations (16). Labored breathing was frequent at ethical endpoints for *BCL6.Mb1.BCL10*^trunc^ animals, and necroscopy revealed high frequency of tumors grossly apparent in lungs, along with splenomegaly and adenopathy (**Fig. 2E-G**). Spleen pathology revealed nodular LBCL tumors in white pulp follicles surrounding central T-cell aggregates (**Fig. 2H**). By IHC, the malignant B cells were positive for B220, CD19, CD20, BCL6, and GFP, with variable Ki67. Nearly three-quarters (72.5%) of *BCL6.Mb1.BCL10*^trunc^ animals developed extranodal tumors involving the lungs (**Fig. 2I**). Lung tumors recapitulated splenic patterns with frequent necrosis. Regions with low necrosis (red boxes) showed malignant cells stained for B220, BCL6, the RSB transgene, and high Ki67 and were surrounded by clusters of T cells (**Fig. 2J**). These results therefore demonstrate lymphogenic synergy between BCL10 GoF and BCL6 over-expression targeted to B cells, recapitulating the genomics of many BN2/C1 cases. These results left unclear the B cell of origin for these tumors, but we noted a consistent lack of GCs by peanut antigen (PNA) staining in spleens (**Fig. 2H**), prompting further investigation.

**Figure 2:**
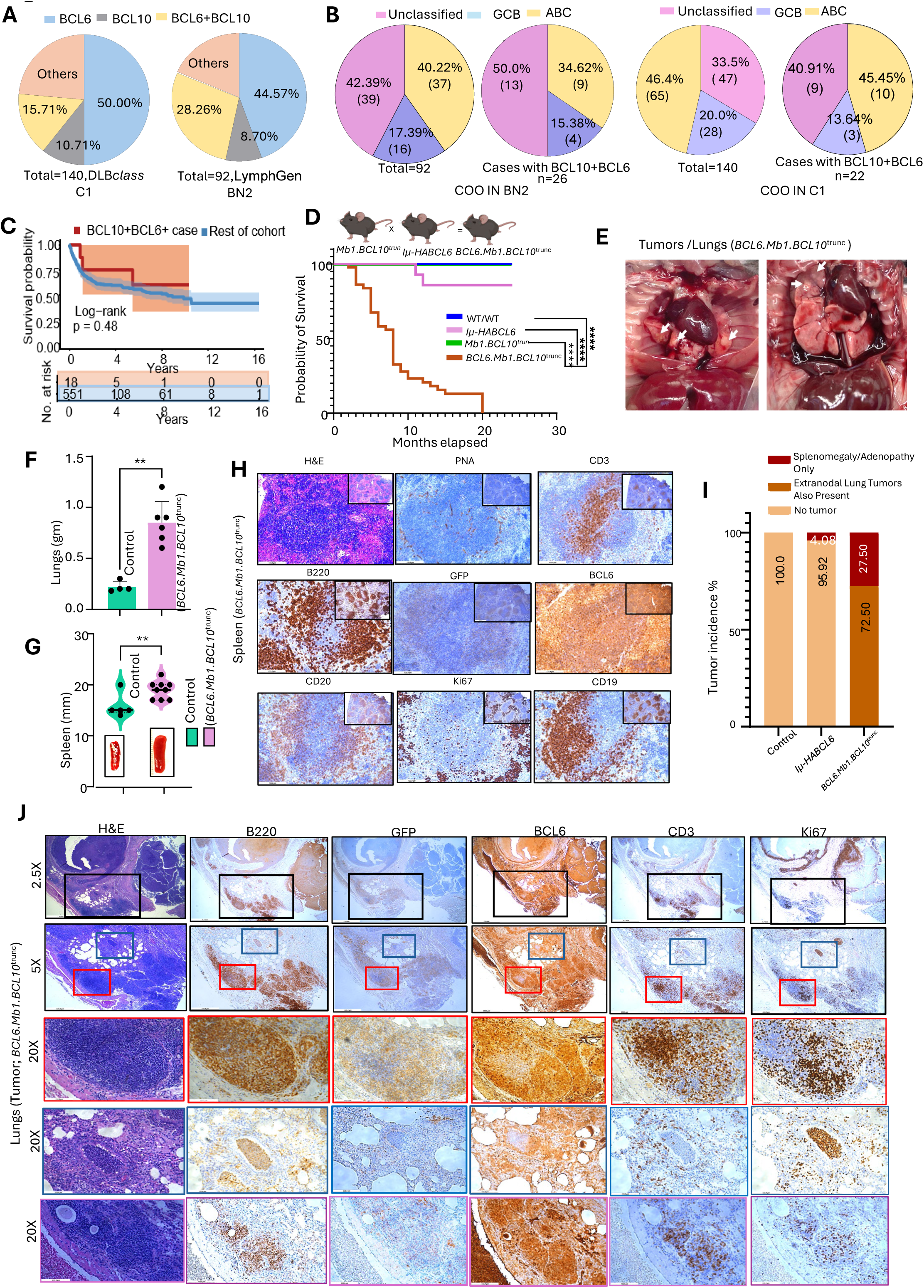
BCL6 Synergizes with BCL10 GoF in Lymphomagenesis. **A**. Pie chart demonstrating distribution of BCL10 and BCL6 mutations among DLBCL patients in BN2/C1 category in DLB*class* and Lymphgen classification. **B**. Distribution based on COO of patients as a whole as well as the BCL10 and BCL6 mutations among the DLB*class* and Lymphgen classification in BN2 clusters and C1 cluster. **C**. Survival probability, all 18 human BCL10+BCL6+ cases vs rest of cohort (n=551) or BN2 group (n=81). Log-rank p = 0.48 (vs rest of cohort) and 0.58 (vs BN2 cases only) using Lymphgen dataset. **D**. Kaplan-Meier **s**urvival curve shows the survival probability for the different genotypes in mice (Mantel-Cox test, P<0.0001. **E**. Anatomy of lungs with tumors (marked in white arrow) in *BCL6.Mb1.BCL10*^trunc^ mice. **F and G**. Weights and size comparison of lungs and spleen respectively among WT/WT and *BCL6.Mb1.BCL10*^trunc^ mice. **H**. Histopathology and immunophenotype of splenic tumors in *BCL6.Mb1.BCL10*^trunc^ mice. Spleen section, stained with hematoxylin and eosin (H&E) or immune-stained for PNA,CD3, B220, GFP, BCL6, CD20, Ki67 and CD19. Insets show broader view of the boxed regions. **I**. Tumor incidence by genotype. Stacked bar graph showing the percentage of mice in each genotype classified as having no tumor (light orange), splenomegaly/adenopathy only (dark red), or splenomegaly/adenopathy with extranodal lung tumors also present (brown) assessed at the endpoint. Control mice remained tumor free (100%), *Iµ-HABCL6* mice developed tumors at low frequency (4.08%), and *BCL6.Mb1.BCL10*^trunc^ showed 100 % tumor incidence, which the majority involved lung tumors (72.05%). **J**. H&E and IHC for tumors (lungs) isolated from *BCL6.Mb1.BCL10*^trunc^ mice. Different tumor lesions with varying degrees of necrosis are shown; lowest necrosis (red boxes), medium necrosis (blue boxes) and highest necrosis (purple markers). Markers; B220, Ki67, CD3, BCL6 and GFP (correlates human BCL10-S136X). Images were acquired on Zeiss microscope equipped with 2.5X objective (scale bars representing 1.1 mm), 5X objective (scale bars representing 535.5 µm), and 20X objective (scale bars representing 133.9 µm).

### *BCL10*^S136X^ imposes a block in germinal center formation not overcome by BCL6

DLBCL derivation from GC and post-GC B cells is well described, but most BN2/C1 cases lack somatic hypermutation (SHM) signatures suggesting non-GC B-cell origin (24,25). Recent combination of the BN2/C1 drivers *NOTCH2* mutation and *SPEN* truncation showed derivation from GC-inexperienced B cells (21). *BCL6* translocations are thought to arise through aberrant AID-mediated recombination in antigen-activated mature B cells, frequently involving immunoglobulin heavy chain (IGH) switch regions (26). Although conventionally associated with GC responses, their precise timing relative to GC entry remains unresolved. It therefore is unclear at what activation stage BCL10 GoF might be most contributory in lymphomagenesis. To assess this, we tested whether intentional GC induction through biweekly immune stimulation with chicken gamma globulin (CGG) would impact BCL10 GoF lymphomagenesis. The inoculated cohort showed *Mb1.BCL10*^trunc^ mice again with no survival difference from littermate genotypes (**Fig. 3A**), though necroscopy revealed mild splenomegaly, with average weights significantly higher than WT (**Fig. 3B**). Histologic examination showed no overt lymphoma in spleens, instead prominent white-pulp follicles that lacked peanut agglutinin (PNA)-positive GCs, in contrast to WT controls (**Fig. 3C-D, Supplementary Fig. 3A**). Since CGG requires priming and adjuvants to mount measurable response, we assessed response to sheep red blood cell (SRBC) inoculation, a well-established inducer of T-dependent GC responses (27). Robust GC formation seen in spleens of WT was nearly absent in *Mb1.BCL10*^trunc^ (**Fig 3E**). To further assess GC responses, we assessed unstimulated animals’ Peyer’s patches (PPs), sites where B cells continuously encounter and are activated by microbial and food antigens. Histology confirmed expression of the transgene in *Mb1.BCL10*^trunc^ PPs in parallel with significantly fewer GC B cells by PNA staining (**Fig. 3F, Supplementary Fig. 3C**). Flow cytometry confirmed reduced GC (FAS^+^GL7^+)^ B cells, and very few of these showed evidence of class switching to IgA (**Fig 3G**). BCL6 regulates GC participation by antigen-stimulated B cells, and GC reactions are correspondingly enhanced in *Iµ-HABCL6* (16). We therefore assessed in premalignant *BCL6.Mb1.BCL10*^trunc^ animals whether BCL6 overcame the GC deficit imposed by BCL10 GoF, but responses to SRBC immunization remained impaired in these animals (**Fig. 3H**). Flow analysis confirmed impaired B-cell activation indicated by retained IgD expression in *BCL6.Mb1.BCL10*^trunc^ **(Fig. 3I)** and diminished FAS^+^IgG1^+^ cells (**Supplementary Fig. 3D**). We next tested whether BCL10 GoF in NB cells imposes a block in CSR, a key step that precedes GC participation during antigen responses (28). We purified splenic NB cells and exposed them to ex vivo immune stimulants that induce CSR (29), replicating in vivo scenarios with and without CD4 T-cell help and/or BCR engagement (LPS, LPS+IL4, LPS+IL4+anti-IgM, anti-CD40L+IL4, or anti-CD40L+IL4+anti-IgM, **Fig 3J**). *Mb1.BCL10*^trunc^ displayed expected IgG isotypes in response to all and indeed were somewhat more responsive than WT to those that did not include BCR cross-linking (**Fig. 3K-P**). When αAnti-IgM was included, effects were either the same as WT (anti-CD40+IL4+anti-IgM) or had fallen behind by 96 hours (LPS+IL4+anti-IgM), suggesting a differential response to BCR engagement (investigated further next section). We also evaluated for defects in BCR clonal diversity, which can impair GC responses in vivo (30) but found no discernible differences (**Supplementary Fig. 3E)**. BCL10 GoF-expressing NB cells therefore are CSR-proficient in response to activating stimuli ex vivo but fail to do so in situ, inhibiting T-cell dependent GC formation, even when BCL6 is concomitantly activated.

**Figure 3:**
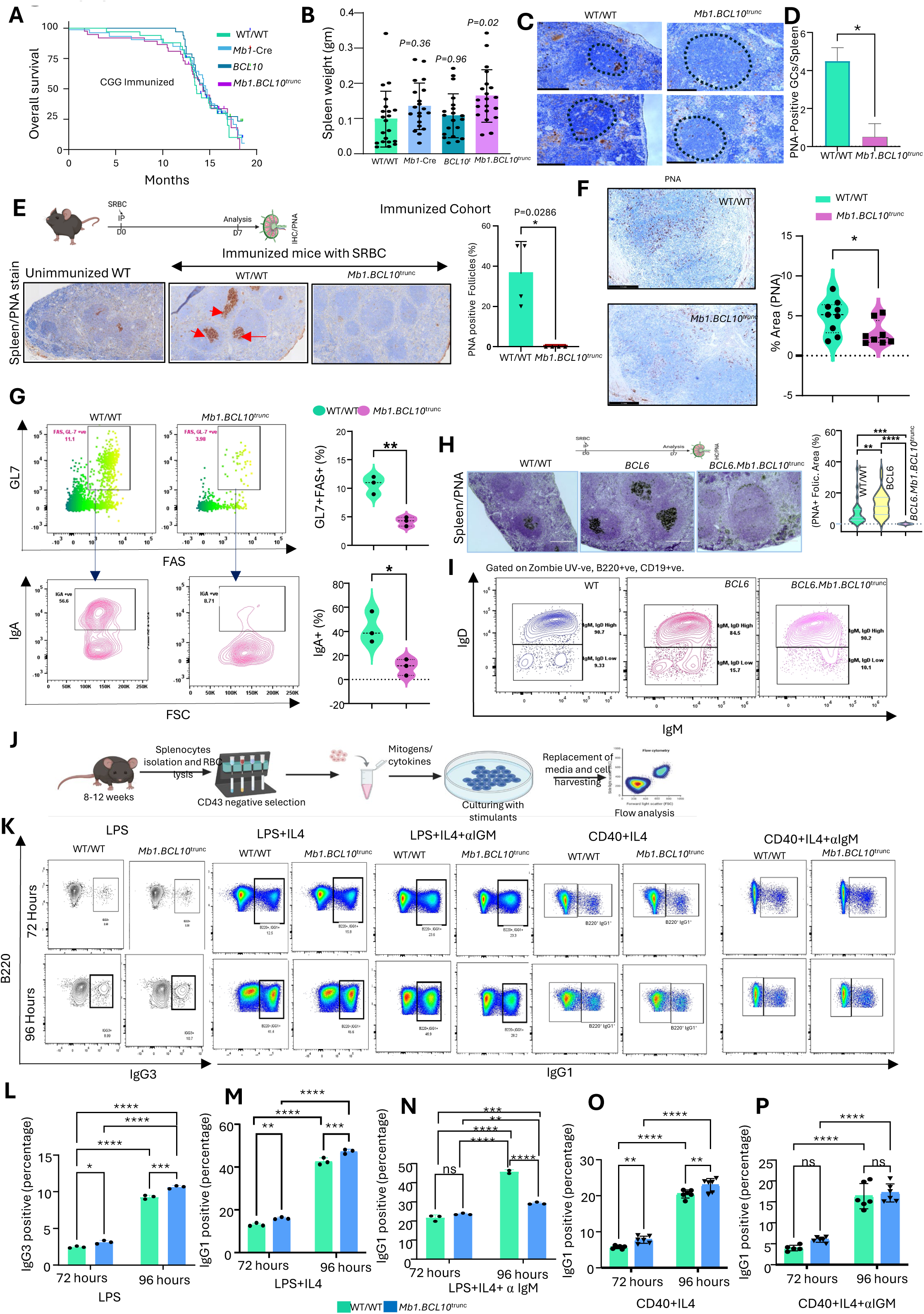
*BCL10*^S136X^ imposes a block in germinal center formation not overcome by BCL6. **A**. Overall survival of CGG immunized mice. Kaplan-Meier survival curves for WT/WT, *Mb1*-cre, *BCL10* and *Mb1.BCL10*^trunc^ following immunization with CGG. Survival curve did not differ significantly between genotypes. **B**. Spleen weights across CGG immunized genotypes at endpoint (n=20 mice per genotype) and p values as indicated in the figure. **C**. Reduced germinal center formation in *Mb1.BCL10*^trunc^ (right) mice immunized with CGG, stained with PNA to identify germinal centers. Two representative fields per genotype are shown. Dashed outlines indicate B cell follicles. WT/WT spleens contained well-formed PNA^+^ GC (brown) within the follicles, whereas follicles in *Mb1.BCL10*^trunc^ were largely devoid of PNA staining. n=3 mice per genotype for PNA staining. **D**. Number of PNA ^+^ GC per spleen section in WT/WT and *Mb1.BCL10*^trunc^. p<0.05. **E**. Schematic of experimental plan (top). PNA stain of SRBC (7-day post immunization) immunized spleens of WT/WT, *Mb1.BCL10*^trunc^. Quantification of percentage PNA positive follicles in spleen sections (right) **F**. Reduced PNA^+^ area in unimmunized *Mb1.BCL10*^trunc^ payer’s patches (PP). Left: representative PP sections from unimmunized WT/WT (top) and *Mb1.BCL10*^trunc^ (bottom). Right: quantification of PNA^+^ staining expressed as percentage of total splenic area. **G**. Representative flow cytometry plots (left panel) and the percentage of GL7^+^, Fas^+^ population (right panel) gated on live B220+, CD19+ population in the SRBC immunized spleen. Top: representative flow cytometry plots of GL7 versus Fas expression on live B220, CD19 positive population isolated from the PP of unimmunized mice. Bottom: IgA expression versus FSC on the cells gated from GL7^+^,FAS^+^ population above (arrows), with gates marking IgA^+^ cells and numbers indicating their frequency. Right: quantification of GL7^+^FAS^+^ GC B cells (top) and IgA^+^ cells (bottom) as a percentage of the parent population. Both populations were significantly reduced in *Mb1.BCL10*^trunc^. **H**. Schematic of experimental plan (top). Left: PNA stain of SRBC (7-day post) immunized spleens of WT/WT, *Iµ-HABCL6, BCL6.Mb1.BCL10*^trunc^. Right: quantification of percentage PNA positive in follicles area in spleen sections (right). **I**. Representative contour plots of IgD versus IgM expression gated on live, B220^+^,CD19^+^ B cells from SRBC immunized spleen of WT/WT, *Iµ-HABCL6, BCL6.Mb1.BCL10*^trunc^ mice. Gates separate IgM/IgD high from IgM/IgD low populations; numbers indicate the frequency of each gate as a percentage of the parent population. **J**. Experimental design for ex vivo stimulation of B cells isolated from the spleen from different genotypes. **K**. Class switching to IgG3 and IgG1 in vitro. Representative flow cytometry plots of B220 versus IgG3 (left two columns) or IgG1 (remaining columns) on cultured B cells from WT/WT and *Mb1.BCL10*^trunc^ at 72 hours (top row) and 96 hours (bottom row). Cells were stimulated with LPS, LPS+IL4, LPS+IL4+anti-IgM, CD40+IL4, or CD40+IL4+anti-IgM as indicated. Gates identify switched B220^+^IgG3^+^ or B220^+^IgG1^+^ cells; numbers indicate the frequency within the parent gate. Representative of either three or five independent experiments. **L-P**. Quantification of class-switch recombination. Percentage of IgG3^+^(L), LPS+IL4 (M), LPS+IL4+anti-IgM (N), CD40+IL4 (O), or CD40+IL4+anti-IgM (P). Results are either three or five independent experiments.

### *BCL10*^trunc^ Promotes Co-Stimulation-Independent Proliferation and Premature Exit from GC Reactions in Premalignant B cells

BCR stimulation of NB cells without reinforcing co-stimulatory signals typically induces anergy or deletion by apoptosis (31,32). Because BCL10 GoF provides CBM activation independently of upstream signals, we hypothesized its contribution to lymphomagenesis could relate to reduced dependence on co-stimulation. In response to the CSR-inducing stimuli employed above (**Fig. 3J-P**), *Mb1.BCL10*^trunc^ splenic NB cells proliferated the same or somewhat more than WT, indicated by increased numbers of cells in higher generation numbers tracked by Cell Trace Violet (CTV) labeling (**Supplementary Fig. 4A-E**). But a major difference emerged in response to BCR cross-linking in isolation. While WT cells as expected were mostly depleted in response to anti-IgM, *Mb1.BCL10*^trunc^ cells survived and proliferated robustly (**Fig. 4A**). Stability of the G0 peak throughout the time course (red arrows) suggests resistance to apoptosis provided by BCL10 GoF, consistent with our previous observations in engineered human cells (5), and an apoptosis assay confirmed significantly fewer apoptotic cells (**Supplementary Fig. 4F**). For further assessment, we performed RNA-seq on pre-stimulated cells and 72 hours post-anti-IgM. Insufficient WT cells survived to 72 hours for analysis, but analysis of *Mb1.BCL10*^trunc^ showed significant induction of proliferative gene-expression and down regulation of apoptosis (**Fig. 4B, Supplementary Fig. 4G**). BCL10 GoF therefore enhances survival and proliferation in response to BCR stimulation alone, substituting at least in part for co-stimulation from T-cell help or TLR engagement that NB cells typically require.

**Figure 4:**
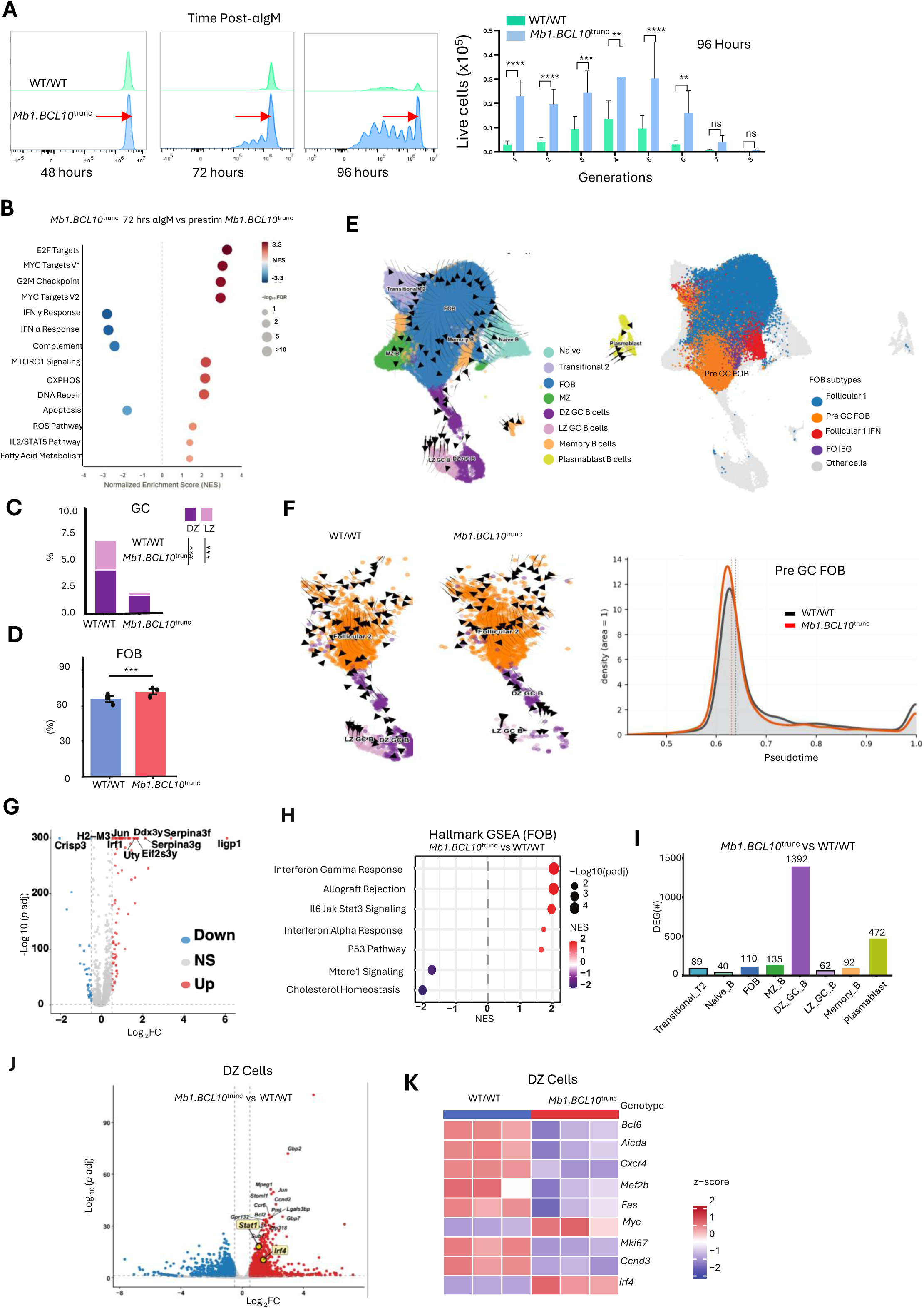
*BCL10*^trunc^ Promotes Co-Stimulation-Independent Proliferation and Premature Exit from GC Reactions in Premalignant B cells. **A**. Enhanced proliferation of *Mb1.BCL10*^trunc^ B cells following BCR stimulation. Left: representative cell trace violet (CTV) dye dilution histograms of purified (CD43-negative selection) B cells from WT/WT (green) and *Mb1.BCL10*^trunc^ (blue) mice at 48, 72, and 96 hours after stimulation with anti-IgM. Red arrows indicate the undivided (generation 0) peak; successive peaks to the left represent sequential cell divisions. Right: number of live cells (×10^5^) in each division generation at 96 hours, quantified from dye dilution profiles. **B**. Gene set enrichment analysis of *Mb1.BCL10*^trunc^ B cells following BCR stimulation. Dot plot of Hallmark gene set significantly enriched in *Mb1.BCL10*^trunc^ cells at 72 hours post anti-IgM stimulation compared with unstimulated (pre-stimulation) *Mb1.BCL10*^trunc^ B cells. Dot color represents NES (red, positive; blue, negative) and dot size represents −log_10_(FDR). Proliferation-associated programs (E2F targets, MYC targets V1 and V2, G2M checkpoint) were strongly enriched following stimulation. N=3 mice per condition. **C**. GC B-cell frequency split into dark zone (DZ) and light zone (LZ) fractions across the WT/WT, *Mb1.BCL10*^trunc^ mice; both fractions are markedly reduced in *Mb1.BCL10*^trunc^ mice (**). These cells are harvested from the spleen after 7 days post SRBC immunization and processed for single cell sequencing as described in supplementary figure 5. Plots represent scRNA-seq derived populations. **D**. Frequency of follicular B cells (FOB); FOBs are significantly increased in *Mb1.BCL10*^trunc^ versus WT/WT in SRBC immunized mice. **E**. UMAP projection of splenic B cells from WT/WT, *Mb1.BCL10*^trunc^, *Iµ-HABCL6*, *BCL6.Mb1.BCL10*^trunc^ and *C*_γ_*1.BCL10*^trunc^, colored by annotated clusters: naïve, Transitional 2, FOB, MZ, DZ GC B cells, LZ GC B cells, Memory B cells and plasmablast B cells. Black arrows show RNA velocity indicating inferred differentiation trajectories. Right: UMAP showing Subclustering of FOB compartment into follicular 1, pre-GC FOB, follicular 1 IFN, and FO IEG subsets; all non-FOB cells are shown in grey. **F**. Pre-GC to germinal center trajectory by genotype. Left and middle: UMAP embeddings of the pre-GC follicular and GC compartments from WT/WT and *Mb1.BCL10*^trunc^ showing follicular 2 (orange), DZ GC B (dark purple), and LZ (light purple) cells. Black arrows show the differentiation trajectory. Right: density distribution of pre-GC FOB cells along pseudotime for WT/WT (black) and *Mb1.BCL10*^trunc^ (red). Dotted vertical lines mark the peak pseudotime value for each genotype. **G.** Volcano plot of differentially expressed genes in FOB cells (*Mb1.BCL10*^trunc^ vs WT/WT; Log2FC vs –Log10 adjusted *P*), with up (red), non-significant (grey) and down-regulated (blue) genes. **H**. Hallmark GSEA of FOB cells (*Mb1.BCL10*^trunc^ vs WT/WT); dot position shows normalized enrichment score (NES), dot size –Log10(adjusted *P*) and color NES. **I**. Number of differentially expressed genes (*Mb1.BCL10*^trunc^ vs WT) per B-cell subset. **J**. Volcano plot of differentially expressed genes in DZ cells (*Mb1.BCL10*^trunc^ vs WT/WT; Log2FC vs –Log10 adjusted *P*), with up (red), non-significant (grey) and down-regulated (blue) genes. IRF4 and STAT1 are marked in bold. **K**. Heatmap (row z-score) of GC-associated genes in *Mb1.BCL10*^trunc^ versus WT in DZ B cells.

We hypothesized that the altered NB response to stimuli plays a role in GC failures underlying lymphomagenesis in *BCL10*^S136X^-expressing B cells. To test this, we assessed cell fates during antigen responses by performing single-cell RNA-sequencing (scRNA-seq) on splenocytes collected seven days after SRBC inoculation. For consistency of analysis throughout this report, we collected samples from n=3 mice per genotype across two control and three experimental genotypes for sequencing as a combined multiplexed run (**Supplementary Fig. 5A**). Results permitted analysis of 199,880 total cells annotated among 10 major cell types (**Supplementary Fig. 5B**). There were 134,273 B cells annotated to eight subtypes (**Supplementary Fig. 5C**), and 43,929 T cells divided to CD4 and CD8 (**Supplementary Fig. 5D**), with these groups separately sub-clustered for analysis in the next section. Comparing WT to *Mb1.BCL10*^trunc^ revealed as expected a significant absence of GC B cells in the latter, impacting especially the light zone (LZ) (**Fig. 4C**). There was instead an increase in follicular B (FOB) cells roughly corresponding, as a percentage of total splenic B cells, to the absent GC cells (**Fig. 4D**). To gain further insight, we reclustered FOB cells to define the specific B cell population undergoing the GC transition (**Fig. 4E**). RNA velocity analysis of these cells further highlighted inability of *Mb1.BCL10*^trunc^ FOB cells to transition to the GC populations (**Fig. 4F**). Our previous work with BCL10 activating mutations showed they promote a cytokine-reinforced gene-expression state marked by IL6, interferon, and other inflammatory signaling activation (5). We hypothesized these same phenomena may occur upon expression in pre-GC B cells. *Mb1.BCL10*^trunc^ FOB cells differentially expressed 110 genes compared to WT (**Fig. 4G**). Gene set enrichment analysis (GSEA) showed consistent with the published results, activation of inflammatory signaling including type I and II interferon and IL6/JAK/STAT3 (**Fig. 4H**). Examining the diminished numbers of *Mb1.BCL10*^trunc^ cells that did progress to GC participation, a near absence of LZ cells suggests arrest and premature exit at the DZ stage. The DZ population in *Mb1.BCL10*^trunc^ differentially expressed 1,392 genes compared to WT, the largest difference of any B-cell subset (**Fig. 4I**), including up-regulation of *Irf4* (**Fig 4J**). IRF4 is induced in late GC cells to promote exit from the reaction (33), and we previously showed it is activated downstream of BCL10 GoF (5). Examination of *Bcl6, Irf4*, and the additional DZ GC markers *Aicda*, *Cxcr4, Mef2b, Fas, Myc, Mki67,* and *Ccnd3* were highly consistent with premature exit from the reaction by the *BCL10*^S136X^-expressing cells (**Fig. 4K**). BCL10 GoF expression in NB cells therefore both inhibited their participation in GCs and promoted IRF4 expression at the DZ stage in parallel with premature exit from the reaction.

### T-Cell Exhaustion is a Prominent Microenvironmental Feature Associated with BCL10 GoF in NB Cells

Having established that B cells expressing mutant BCL10 are not inherently defective in class switching, we hypothesized cell-extrinsic factors, i.e., altered microenvironments, contribute to GC failures. We therefore compared scRNA-seq results (**Supplementary Fig. 5**) among WT, *Iµ-HABCL6*, *Mb1.BCL10*^trunc^, and *BCL6.Mb1.BCL10*^trunc^. There were no significant differences comparing relative proportions of CD4 and CD8 cells (**Supplementary Fig. 5C**), and we reclustered these groups individually (**Supplementary Fig. 6A-E**) (34). We observed that both *BCL10*^trunc^ genotypes were significantly more infiltrated by exhausted (Tex) and less infiltrated by naïve (Tn) T cells, with CD8 cells most affected (**Fig. 5A-D**). Consistent with the proportional changes in these populations, RNA velocity analysis revealed a differentiation trajectory from Tn to Tex that was prominent in both *BCL10*^trunc^ genotypes (**Fig. 5E-F**). These expanded proportions of TOX-expressing CD8 T cells were unexpected. TOX is canonically induced by chronic T cell receptor engagement and sustained NFAT activation (35), and is therefore characteristic of persistent antigenic stimulation. This biology is not consistent with a change likely to have developed as a consequence of SRBC stimulation after seven days but rather a pre-existing alteration in the microenvironments of the *BCL10*^trunc^ genotypes driven by an altered cytokine milieu. To test this, we examined samples from unstimulated animals, confirming both *BCL10*^trunc^ genotypes have increased Tex cells prior to SRBC inoculation (**Fig. 5G**). Next, we examined intercellular interactions in the single-cell data by CellChat (36). This revealed consistent, significantly stronger B-to-T cell than T-to-B cell interactions most pronounced with *BCL10*^trunc^ and to a lesser extent in *Iµ-HABCL6* (**Fig. 5H**). The strongest B-to-T enrichment targeted CD8^+^ T central memory (Tcm), T effector memory (Tem), T differentiated effector memory (Temra), Tex and T IFN-response (Tisg) subsets, indicating that B cells are the predominant source of altered intracellular signals. (**Supplementary Fig. 6F-G**). GSEA comparing the transgenic genotypes to WT revealed, consistent with the exhausted phenotypes, prominent decline of proliferative gene sets in *Mb1.BCL10*^trunc^ and *BCL6.Mb1.BCL10*^trunc^, an effect largely absents in *Iµ-HABCL6* (**Supplementary Fig. 6I**). Effects again were strongest in CD8 populations, with the *BCL10*^trunc^ genotypes more separated from WT than *Iµ-HABCL6* (**Fig. 5I**). Examination of specific genes showed CD8 populations in both *BCL10*^trunc^ genotypes, in contrast to *Iµ-HABCL6*, showed elevated *Ifng, Hvacr2, Lag3, Pdcd1, Gzmk, Gzmb, Ccl4* and *Prf1* (**Fig. 5J**), with similar patterns also in the CD4 populations (**Supplementary Fig. 6J**). Inferred Tfh-to-B cell interactions were largely preserved in the *BCL10*^trunc^ expressing genotypes relative to WT/WT and *Iµ-HABCL6* controls, arguing against impaired Tfh help as primary driver of the abrogated GCs (**Supplementary Fig. 6H**). Collectively, these genes define a program characteristic of sustained T cell activation within an interferon-rich inflammatory milieu. Concomitant expression of *Pdcd1*, *Lag3*, and *Havcr2* is a canonical signature of persistent activation, and the number of co-expressed inhibitory receptors scales with the degree of functional impairment (35,37). *Gzmk*^+^ CD8+ T cells in particular have been described as a chemokine-producing, inflammation-associated population distinct from conventional cytotoxic effectors, accumulating in chronically inflamed tissue with age (38). Confinement of these patterns to the *BCL10*^trunc^ harboring genotypes indicates it’s a specific consequence of B-cell intrinsic, mutant BCL10-driven signaling rather than a general feature of T-dependent B cell stimulation.

**Figure 5:**
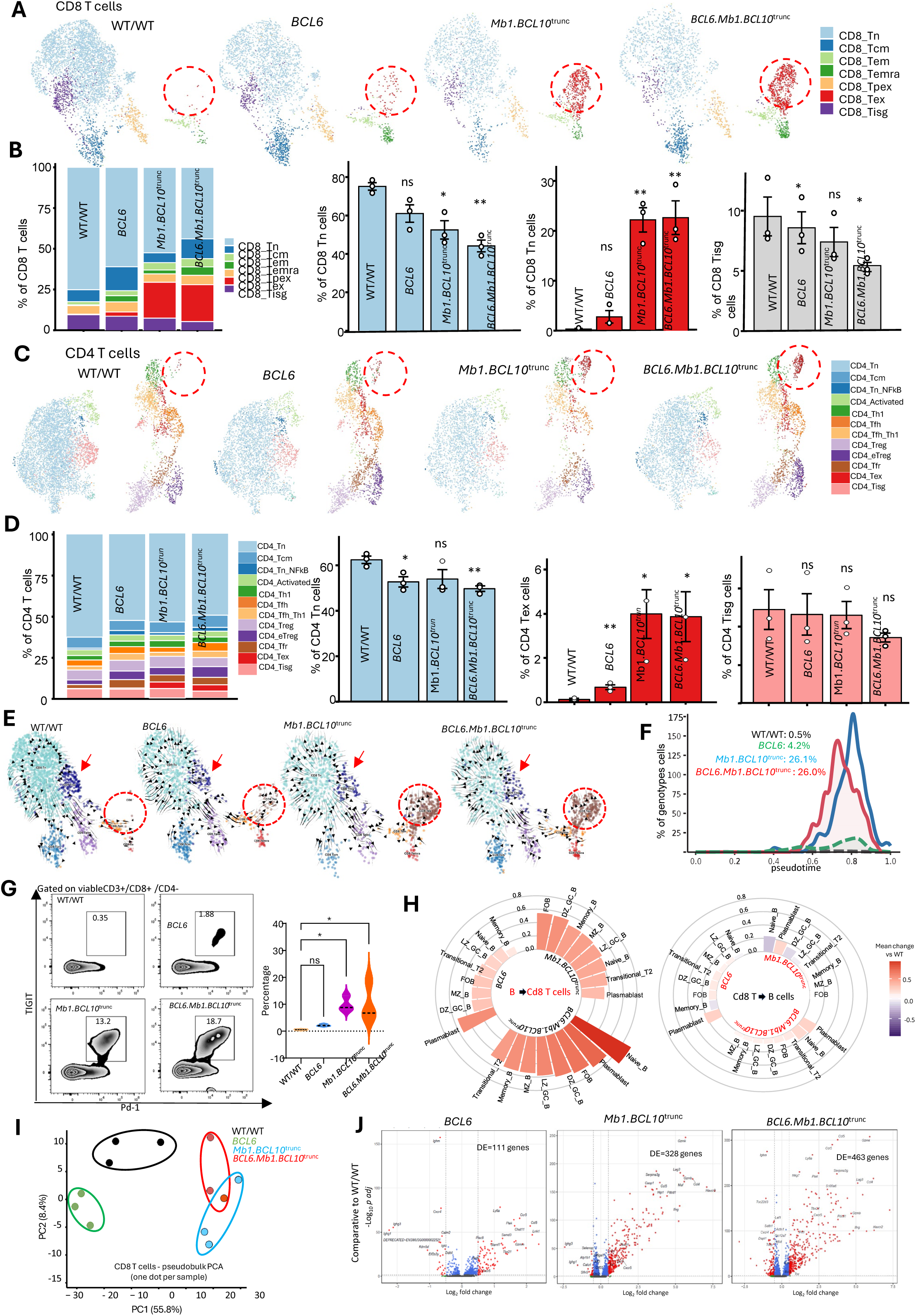
T-Cell Exhaustion is a Prominent Microenvironmental Feature Associated with BCL10 GoF in NB Cells. **A.** CD8 T cell landscape across genotypes. UMAP projections of splenic CD8 T cells from WT/WT, *Mb1.BCL10*^trunc^, *Iµ-HABCL6* and *BCL6.Mb1.BCL10*^trunc^ mice colored by annotated subsets: naïve (Tn), central memory (Tcm), effector memory (Tem), terminally differentiated effector memory (Temra), progenitor exhausted (Tpex), exhausted (Tex), and IFN response stimulated (Tisg). Red dashed circles highlight the exhausted (Tex) compartment, which expanded progressively in *BCL10*^trunc^ mutant genotypes. **B**. Quantification of CD8 subsets. Left: stacked bars showing the proportional composition of CD8 subsets per genotype. Right: Comparison of the Tn, Tex and Tisg populations. Comparison of other populations is mentioned in supplementary figure 6. **C**. CD4 T cell landscape across genotypes. UMAP projections of splenic CD4 T cells from WT/WT, *Mb1.BCL10*^trunc^, *Iµ-HABCL6* and *BCL6.Mb1.BCL10*^trunc^ mice colored by annotated subsets: Tn, Tcm, Tn_NFkB, activated, Th1, Tfh, Tfh_Th1, Treg, effector Treg, Tfr, Tex and Tisg. Red dashed circles highlight the expansion of Tex population among *Mb1.BCL10*^trunc^ and *BCL6.Mb1.BCL10*^trunc^ genotype. **D**. Quantification of CD4 subsets. Left: stacked bars showing the proportional composition of CD4 subsets per genotype. Right: Comparison of the Tn, Tex and Tisg populations. Comparison of other populations is mentioned in supplementary figure 6. **E**. CD8 T cells differentiation trajectories across genotypes. UMAP projections of splenic CD8 T cells from WT/WT, *Mb1.BCL10*^trunc^, *Iµ-HABCL6* and *BCL6.Mb1.BCL10*^trunc^, colored by subset as mentioned above with black arrows showing RNA velocity vectors. Red arrows indicate the transition of cells into Tex; red dashed circles highlight the exhausted T cell compartment, which was minimal in WT/WT and markedly expanded in *Mb1.BCL10*^trunc^ and *BCL6.Mb1.BCL10*^trunc^ genotypes. **F**. Pseudotime distribution of CD8 T cells by genotype. Density of CD8 T cells along pseudotime for each genotype. Both *BCL10*^trunc^ genotypes showed a pronounced accumulation of cells at early pseudotime relative to WT/WT and *Iµ-HABCL6* controls. **G**. TIGIT and PD-1 co-expression on CD8 T cells in unimmunized mice. Left: representative contour plots of TIGIT versus PD-1 on viable CD3^+^CD8^+^and CD4^−^splenic T cells from each genotype. Right: quantification across genotypes. Each dot represents a mouse. n=3 mice per genotype. *BCL10*^trunc^ genotypes have high percentage of TIGIT and PD-1 expressing CD8 T cells. **H**. Intercellular signaling between B cells and CD8 T cells. Circular bar plots showing the mean change relative to WT/WT in predicted interaction strength for B cell towards CD8 T cell signaling (left) and CD8 T cell towards B cell signaling (right), across *Mb1.BCL10*^trunc^, *Iµ-HABCL6* and *BCL6.Mb1.BCL10*^trunc^ genotypes (concentric rings). Each bar represents one B cell subset. Bar height and color indicate the magnitude and direction of change (red, increases; purple, decreased). Analysis performed with CellChat. **I**. PCA analysis of pseudobulk expression profiles from splenic CD8 T cells, with one point per mouse (n=3 mice per genotype): WT/WT (black), *Iµ-HABCL6* (green)*, Mb1.BCL10*^trunc^ (blue) and *BCL6.Mb1.BCL10*^trunc^ (red). Ellipses are drawn to indicate genotype groupings. Both *BCL10*^trunc^ genotypes are clustered together and separated from WT/WT and *Iµ-HABCL6* genotype. **J**. Volcano plots of differentially expressed genes in CD8 T cells from *Iµ-HABCL6, Mb1.BCL10*^trunc^ and *BCL6.Mb1.BCL10*^trunc^.

We also assessed if failure of tumors to develop in primary *Mb1.BCL10*^trunc^ mice might be due to these microenvironmental differences with WT. We transplanted *BCL10*^trunc^ tumor cells (**Fig. 1G**) to 14 WT and 14 *Mb1.BCL10*^trunc^ animals (1 x 10^6^ per recipient), with five animals for each recipient genotype receiving 3.5 Gy preparative lymphodepleting radiation before injection while the rest were non-radiated (**Supplementary Fig. 6K**). The radiation as expected accelerated onset of fatal lymphomas, but there was no significant difference in tumor onset between the recipient genotypes under either condition (**Supplementary Fig. 6L-M)**. Histology from the non-irradiated condition revealed diffuse large-cell infiltration in WT recipients as in the prior experiments, while *Mb1.BCL10*^trunc^ recipients exhibited nodular patterns with discrete aggregates of large tumor cells (**Supplementary Fig. 6N)**. The WT tumors had areas of necrosis consistent with more unrestrained proliferation. Nodularity of the *Mb1.BCL10*^trunc^ host tumor-involved spleens compared to WT was evident also grossly (**Supplementary Fig. 6O)**. Less permissive microenvironments in *Mb1.BCL10*^trunc^ animals may therefore have inhibited the outgrowth of lymphomatous clones in primary animals, but differences were not sufficient to result in survival differences for secondary recipients, with or without lymphodepletion.

### A T-bet^+^/IL9R Extrafollicular Memory B-Cell Phenotype Underlies BCL6-BCL10 GoF Lymphomagenesis

Mutant-BCL10 activity in NB cells, with or without concomitant BCL6, promoted GC failure and accumulation of exhausted T cells in splenic microenvironments, but only the combination led to lymphoma onset in primary animals. The synergistic interaction therefore occurred in a non-GC population, possibly in cells that underwent extrafollicular (EF) antigen responses. To investigate this, we compared B-cell gene expression patterns among WT, *Iµ-HABCL6, Mb1.BCL10*^trunc^, and *BCL6.Mb1.BCL10*^trunc^. PCA from the single-cell data (**Supplementary Fig. 5**) showed unique clustering of the triple-heterozygous genotype, driven by 125 unique DEGs relative to WT, compared to 48 and 46 for *Mb1.BCL10*^trunc^ and *Iµ-HABCL6* respectively (**Fig. 6A-B**). Directly comparing *Mb1.BCL10*^trunc^ with or without *BCL6* by GSEA showed multiple pathways upregulated in *BCL6.Mb1.BCL10*^trunc^ B cells, including proliferative and metabolic processes (G2/M Checkpoint, E2F Targets, Oxidative Phosphorylation, **Fig. 6C**). To see if this corresponds to the biology of human tumors, we examined gene-expression data from LymphGen clinical cases (3,4). Cases with both *BCL10* GoF and *BCL6* translocation compared with the rest of the cohort revealed substantial overlap with the pathways identified in *BCL6.Mb1.BCL10*^trunc^ (**Fig. 6D-E**). In both the human and murine comparisons IL6/JAK/STAT3 signaling was prominent, including being further enhanced in *BCL6.Mb1.BCL10*^trunc^ over *Mb1.BCL10*^trunc^, already up-regulated vs WT (**Fig. 4H**). Inflammatory and interferon signaling, by contrast, were significantly reduced commonly in both mouse and human GSEA. The combination of BCL6 deregulation and BCL10 GoF therefore promotes gene-expression changes with multiple oncogenic stigmata, including cell cycle progression and reduced intracellular inflammatory responses. These stigmata are consistent with BCL6 activation, which promotes both enhanced proliferation and dampening of intracellular responses to processes such as active DNA damage (8,14). However, *BCL6* alone did not replicate transformation rapidity or penetrance of the *BCL6.Mb1.BCL10*^trunc^ genotype, so we investigated further the basis for synergy. Examining specific up-regulated genes in *BCL6.Mb1.BCL10*^trunc^ compared to *Mb1.BCL10*^trunc^, we note as expected *Bcl6* and also *Il9r*, encoding interleukin-9 receptor (IL9R), which promotes STAT3 activation and DLBCL cell survival (**Fig. 6F**) (39–41). STAT3 signaling was specifically up-regulated in *BCL6.Mb1.BCL10*^trunc^ above the *Mb1.BCL10*^trunc^ genotype (**Fig. 6C**), and we note that activated STAT3 directly binds to and up-regulates expression of *BCL6* (42). Therefore, we hypothesized BCL10 GoF promotes further enhancement of *Bcl6* expression compared to the *BCL6* transgene alone through cytokine receptor-mediated STAT3 activation. Comparison of *Il9r* and *Bcl6* mRNA levels between genotypes confirmed *BCL6.Mb1.BCL10*^trunc^ cells had the highest levels of both (**Fig. 6G-H**).

**Figure 6:**
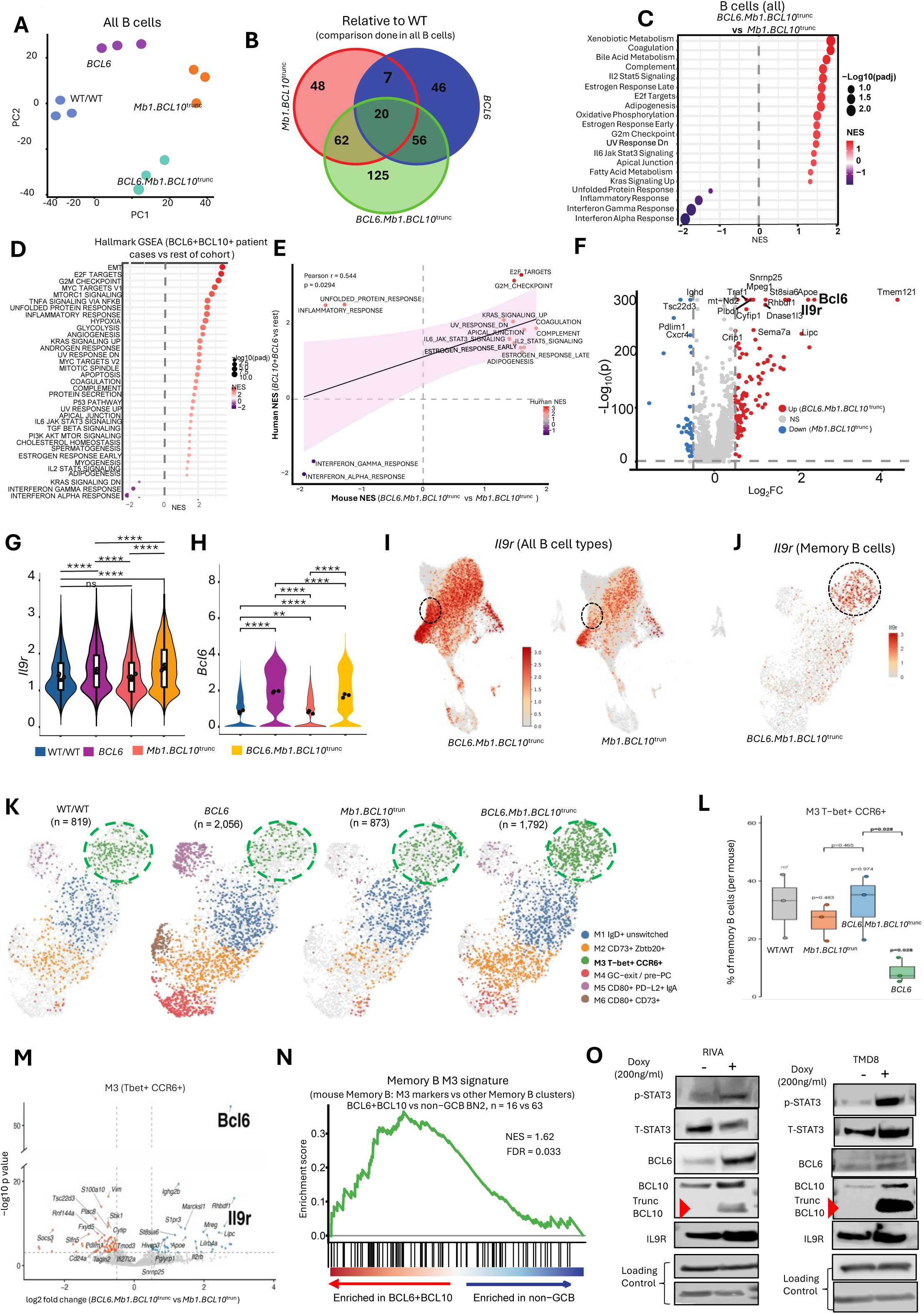
A T-bet^+^/IL9R Extrafollicular Memory B-Cell Phenotype Underlies BCL6-BCL10 GoF Lymphomagenesis. **A**. Principal-component analysis (PC1 vs PC2) of the B-cell compartment, separating the four genotypes (WT, *Mb1.BCL10*^trunc^, *Iµ-HABCL6*, *BCL6.Mb1.BCL10*^trunc^). **B**. Venn diagram of differentially expressed genes shared among the *Mb1.BCL10*^trunc^, *Iµ-HABCL6* and *BCL6.Mb1.BCL10*^trunc^ comparisons (versus WT): 48 genes unique to *Mb1.BCL10*^trunc^, 46 to *Iµ-HABCL6* and 125 to TH; pairwise overlaps of 7 (*Mb1.BCL10*^trunc^ vs *Iµ-HABCL6*), 62 (*Mb1.BCL10*^trunc^ vs *BCL6.Mb1.BCL10*^trunc^) and 56 (*Iµ-HABCL6* vs *BCL6.Mb1.BCL10*^trunc^), with 20 genes common to all three. **C**. Hallmark GSEA of the total B-cell compartment (*BCL6.Mb1.BCL10*^trunc^ vs *Mb1.BCL10*^trunc^); dot position indicates normalized enrichment score (NES), size –Log10 (adjusted *P*) and color NES. Metabolic and proliferative programmes (xenobiotic metabolism, coagulation, bile-acid metabolism, complement, IL2/STAT5 signaling, E2F targets, oxidative phosphorylation, G2M checkpoint, IL6/JAK/STAT3 signaling) are positively enriched, whereas interferon-alpha/gamma responses, inflammatory response and the unfolded-protein response are negatively enriched. **D**. Hallmark GSEA, human: 18 BCL10+BCL6+ DLBCL cases vs rest of cohort. Dot color = NES, size = −log10 (padj), padj < 0.05. **E**. Correlation of Hallmark NES between human (A) and mouse (B), shared pathways. Pearson r = 0.544, p = 0.029, n=16. NF-κB/proliferation pathways concordant (upper right); interferon response concordant (lower left). **F**. Volcano plot of differentially expressed genes (*BCL6.Mb1.BCL10*^trunc^ vs *Mb1.BCL10*^trunc^; Log2FC vs –Log10 *P*) with up-regulated in *BCL6.Mb1.BCL10*^trunc^ (red), non-significant (grey) and down-regulated in *Mb1.BCL10*^trunc^ (blue). **G.** Violin plot of *Il9r* expression (log) in FOB cells across WT, *Mb1.BCL10*^trunc^, *Iµ-HABCL6* and *BCL6.Mb1.BCL10*^trunc^; expression is comparable between WT and *Mb1.BCL10*^trunc^ (ns) and significantly altered in the other comparisons (****). **H**. Violin plot of *Bcl6* expression across WT/WT, *Iµ-HABCL6*, *Mb1.BCL10*^trunc^ and *BCL6.Mb1.BCL10*^trunc^; *Bcl6* is significantly elevated in the *Iµ-HABCL6* and *BCL6.Mb1.BCL10*^trunc^ genotypes (****, **). **I**. UMAP projections of splenic B cells from *BCL6.Mb1.BCL10*^trunc^ (left) and *Mb1.BCL10*^trunc^ (right) mice, colored by normalized *Il9r* expression (scale as shown). Dashed circles indicate the memory B cells cluster with markedly enriched *Il9r* expression in *BCL6.Mb1.BCL10*^trunc^ compared to *Mb1.BCL10*^trunc^. **J**. UMAP embedding of splenic B cells from *BCL6.Mb1.BCL10*^trunc^ mice colored by normalized *Il9r* expression, with the dashed circle marking the memory B cell cluster, which showed enriched *Il9r* expression relative to other subsets. **K**. Memory B cell subset composition across the genotypes. UMAP projections of splenic memory B cells from WT, *Mb1.BCL10*^trunc^, *Iµ-HABCL6* and *BCL6.Mb1.BCL10*^trunc^ colored by memory B cell subset. Cell numbers per genotype are indicated. Green dashed circles highlight the M3 Tbet^+^CCR6^+^ cells among total memory B cells. **L**. Percentage of M3 Tbet^+^CCR6^+^ cells among total memory B cells, calculated per mouse. WT/WT was taken as reference to calculate the p value. **M**. Volcano plot of differentially expressed genes (*BCL6.Mb1.BCL10*^trunc^ vs *Mb1.BCL10*^trunc^; Log2FC vs –Log10 *P*) in M3 subset with up-regulated in *BCL6.Mb1.BCL10*^trunc^ (blue), non-significant (grey) and down-regulated in *BCL6.Mb1.BCL10*^trunc^ (red). *BCl6* and *IL9r* were highlighted in bold. **N.** GSEA of the mouse M3 Tbet^+^CCR6^+^ memory B cell signature (defined as genes differentially expressed in M3 versus all other memory B clusters, panel K) in human DLBCL cases with concurrent *BCL6* and *BCL10* alterations (n=16) compared with non-GCB/BN2 cases (n=63) from. **O**. Western blot showing different markers (p-stat3, BCL6, truncated BCL10, IL9r and total STAT3) in 24 hours doxycycline induced RIVA and TMD8 cell lines.

We hypothesized gene expression patterns specific to the *BCL6.Mb1.BCL10*^trunc^ genotype could help reveal the underlying non-GC/EF cellular state most responsible for the synergistic interaction. We therefore projected *Il9r* across B-cell subsets and found particular enhancement in the memory B cell (MBC) population of *BCL6.Mb1.BCL10*^trunc^ animals (**Fig. 6I**). Reclustered MBCs showed a specific subgroup driving the elevated expression (**Fig. 6J**), which annotation revealed as M3 (T-bet^+^ CCR6^+^) MBCs (**Fig. 6K, Supplementary Fig. 7A-D**). T-bet memory populations arise predominantly outside GCs and participate in IL9R-dependent EF-like memory responses (43,44). Expansion of this population was a direct consequence of *BCL6* expression, representing a significant change when added to *BCL10*^trunc^ expression alone (**Fig. 6L**). Moreover, these cells in the *BCL6.Mb1.BCL10*^trunc^ genotype with both oncogenes working together had much higher expression of *Il9r* (**Fig. 6M**). These results suggest BCL10/BCL6 co-deregulation establishes an IL9R-high extrafollicular memory-like B-cell state that serves as a reservoir for malignant transformation. To test if this plays out in clinical DLBCL, we further analyzed the LymphGen dataset. Assessing cases with co-occurrence of *BCL10* mutation and *BCL6* translocation compared to non-GCB cases as a whole showed significant enrichment of the M3 MBC gene-expression signature (**Fig. 6N**). Finally, we tested functionally in DLBCL cell lines the effect of adding *BCL10*^trunc^ to baseline *BCL6* deregulation using systems we previously engineered with tet-on *BCL10*^S136X^ (5). RIVA cells are a model for BN2/C1 cases with both a *BCL6* translocation and *NOTCH2* mutation (1,4). Induction of *BCL10*^trunc^ in these cells induced expression of IL9R, promoted STAT3 activation, and significantly enhanced BCL6 expression (**Fig. 6O**). Results were the same in TMD8, an independent baseline *BCL6*-expressing system (**Fig. 6P**). These data suggest BCL10 GoF collaborates with BCL6 deregulation in DLBCL through BCL10-driven STAT3 activation that further enhances BCL6 expression to promote malignant transformation. Distinctive M3 expansion with elevated IL9R in a GC-deficient background underlies these effects with their gene-expression patterns carrying through to corresponding clinical cases.

### BCL6/BCL10 GoF Synergy Favors GC-Independent Lymphomagenesis Even in GC-Experienced B Cells

Our results establish oncogenic synergy between BCL10 GoF and BCL6 promoting lymphomas that arise from GC-inexperienced B cells when the genes are co-expressed under lineage-specific promoters active prior to GC entry, Mb1 and Iµ respectively. Timing of acquisition of BCL10 mutations in clinical cases relative to stage of B-cell development or activation, however, is not clear in published genomics data. To understand if the expression of BCL10 mutations at later stage of BCL10 activation can lead to different phenotype, we crossed *RSB* to *C*_γ_*1Cre*, creating *C*_γ_*1.BCL10*^trunc^ animals, to induce expression of the transgene at the point of CSR (45). We first assessed GC responses in these animals through SRBC inoculation and found robust formation by PNA staining enhanced compared to WT, in marked contrast to the results in *Mb1.BCL10*^trunc^ (**Fig. 7A**). Flow analysis of splenic B cells confirmed increase of activated (IgD^−^,IgM^+^) cells, accumulation of GC (FAS^+^GL7^+^) cells, and successful class switching to IgG1 in *C*_γ_*1.BCL10*^trunc^, effects largely absent in *Mb1.BCL10*^trunc^ (**Supplementary Fig. 8A-C**). We also confirmed, as expected, that *C*_γ_*1.BCL10*^trunc^ NB cells, in which the transgene is not yet activated, do not have enhanced proliferation in response to anti-IgM without co-stimulation (**Supplementary Fig. 8D**). ScRNA-seq results (**Supplementary Fig. 5**) for *C*_γ_*1.BCL10*^trunc^ confirmed transgene expression limited to GC and post-GC populations and robust presence of DZ and LZ cells, indicating that the later induction of BCL10 GoF did not promote premature GC exit, in contrast to *Mb1.BCL10*^trunc^ (**Fig. 4K**, **Fig. 7B-C**). Splenic microenvironments also were significantly less infiltrated by Tex cells **(Fig. 7D)**. Since *C*_γ_*1*-driven expression of BCL10 GoF promoted hyperproliferative GCs, we assessed for tumor onset in *C*_γ_*1.BCL10*^trunc^ animals subjected SRBC inoculation every 30 days. Like *Mb1.BCL10*^trunc^ animals, however, the genotype did not result in lymphoma onset in animals followed out to 22 months (**Fig 7E**). We assessed splenocytes after six SRBC inoculations and found that despite the hyperproliferative response to initial inoculation, *C*_γ_*1.BCL10*^trunc^ lost later GC responses in contrast to control *C*_γ_*1-Cre* animals (**Fig. 7F**). Flow cytometry showed GPF^+^ B cells in spleens, confirming the presence of GC-experienced cells from past immunizations, but impaired induction of active GC (FAS^+^/GL7^+^) cells (**Supplementary Fig. 8E**). Therefore, *BCL10*^trunc^ induction directly in active GCs did not lead to lymphoma onset, and accumulation in spleens of post-GC cells expressing the transgene over time impaired subsequent GC responses. For further assessment, we crossed the *C*_γ_*1.BCL10*^trunc^ animals with *Iµ-HABCL6* to generate triple heterozygous *BCL6.C*_γ_*1.BCL10*^trunc^, which also were subjected to repeated SRBC immunization to promote Cre expression. These began dying after 4 months of repeated immunization (**Fig. 7E**). Necroscopy showed a lung tumor phenotype similar to that observed in *BCL6.Mb1.BCL10*^trunc^, along with splenomegaly, LBCL tumors disrupting splenic architecture, and persistent absence of GC cells (**Figure 7G-J**). Thus, *BCL6* overexpression combined with *BCL10* GoF favors transformation of non-GC populations, even in post GC B cells provided regular stimulus for GC re-entry.

**Figure 7:**
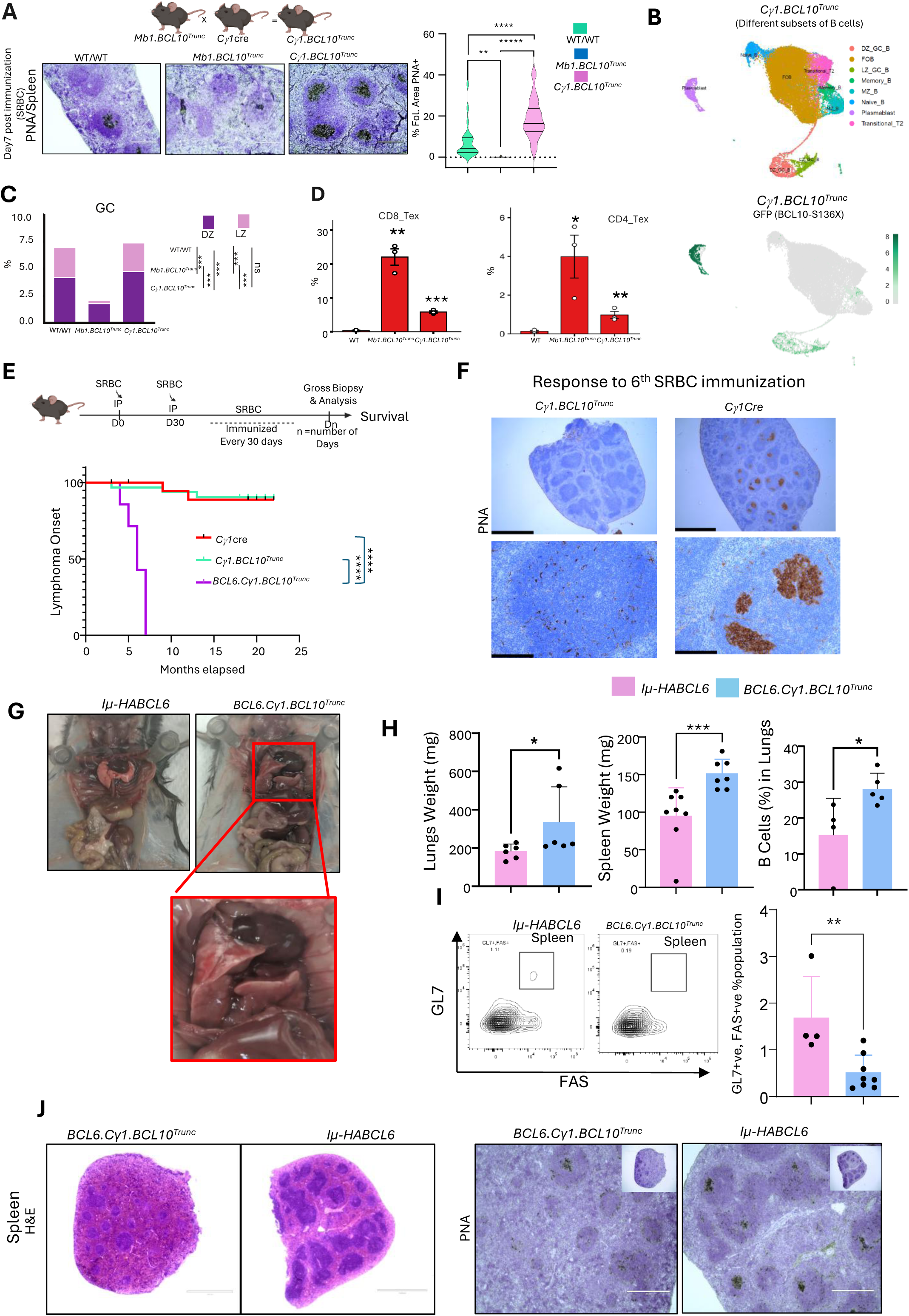
BCL6/BCL10 GoF Synergy Favors GC-Independent Lymphomagenesis Even in GC-Experienced B Cells. **A.** Experimental design and PNA for spleen isolated from mice 7 days post SRBC immunization and quantitative bar plots showing the PNA positive area per follicle (n=4) across WT/WT, *Mb1.BCL10*^trunc^ and *C*_γ_*1.BCL10*^trunc^ **B.** B cell subsets and *BCL10*^trunc^ expression in *C*_γ_*1.BCL10*^trunc^. Top: UMAP projections of splenic B cells from *C*_γ_*1.BCL10*^trunc^ mice, colored by annotated subset. Bottom: the same UMAP projections colored by GFP expression, indicating expression of the mutant BCL10 following Cre-mediated excision of the STOP cassette. GFP expression was confined to GC, plasmablasts and memory B cells populations consistent with the GC restricted activity of *C*_γ_*1* promoter expressing Cre recombinase. **C**. GC B-cell frequency split into DZ and LZ fractions across the WT/WT, *Mb1.BCL10*^trunc^ and *C*_γ_*1.BCL10*^trunc^ mice; both fractions are markedly reduced in *Mb1.BCL10*^trunc^ mice (***) and enhanced in *C*_γ_*1.BCL10* mice (***). This data is the read out of Sc-RNA seq data mention in supplementary figure 5. **D**. Percentage of CD8 and CD4 T exhausted cells across SRBC immunized WT/WT, *Mb1.BCL10*^trunc^ and *C*_γ_*1.BCL10*^trunc^ mice. Read out of supplementary figure 5. **E.** Top: schematic presentation of the experiment for SRBC repeated immunizations in *C*_γ_*1.BCL10*^trunc^ and *BCL6.C*_γ_*1.BCL10*^trunc^ mice. Bottom: Kaplan-Meier **s**urvival curve shows the survival probability across *C*_γ_*1*cre, *Mb1.BCL10*^trunc^, *C*_γ_*1.BCL10*^trunc^ and *BCL6.C*_γ_*1.BCL10*^trunc^ genotypes in mice (Mantel-Cox test, P<0.0001). **F**. Representative PNA staining for *C*_γ_*1.BCL10*^trunc^ and *C*_γ_*1-Cre* showing reduced PNA staining in the *C*_γ_*1.BCL10*^trunc^ compared to either *C*_γ_*1-Cre* after repeated SRBC immunization or compared to *C*_γ_*1.BCL10*^trunc^ itself after single SRBC immunization. **G**. Gross necropsy showing tumors in lungs in the *BCL6.C*_γ_*1.BCL10*^trunc^ mice (marked in red box). **H**. Bar graphs showing the lungs and spleen weight (mg) and percentage of B cells in lungs in *BCL6.C*_γ_*1.BCL10*^trunc^ versus *Iµ-HABCL6*. **I**. Representative flow cytometry plots showing *BCL6.C*_γ_*1.BCL10*^trunc^ deficit in Fas^+^GL7^+^ B cells gated on B220+, CD19+ viable B cells comparing to *Iµ-HABCL6* collected at the endpoint after repeated SRBC immunization. **J**. Left: H&E for the spleen sections harvested from repeatedly SRBC immunized mice across *BCL6.C*_γ_*1.BCL10*^trunc^ and *Iµ-HABCL6.* Right: PNA staining for GCs showing reduced PNA^+^ cells in *BCL6.C*_γ_*1.BCL10*^trunc^ mice comparing to *Iµ-HABCL6* control.

## Discussion

Gene expression-based COO subclassification of DLBCL at the dawn of the genomics era provided the first glimpses of the profound heterogeneity underlying pathogenesis of the most common lymphoma diagnosis (46–48). Current classifications have evolved ABC, GCB, and unclassified COO groupings to 5-7 genomic clusters based on recurrent single-gene mutations and segment-level chromosomal copy-number gains and losses (1–4). Even within individual clusters, however, evidence for significant heterogeneity of disease-driving alterations is prominent. BN2/C1 cases provide an illustrative example. *BCL6* translocations and *NOTCH2* activating mutations are defining, but cases may have only one or even neither of these, and all have additional drivers present. Therefore, precision genetic modeling in mice provides opportunity to define the basis for transformation in specific contexts. Here, we started with BCL10 GoF which alone was able to promote DLBCL only after selection ex vivo for B-cell clones with malignant potential upon adoptive transfer to secondary recipients (**Fig. 1**). Addition of *BCL6* deregulation, however, replicating a co-occurrence frequent in BN2/C1 cases, promoted tumor onset in 100% of primary animals (**Fig. 2**).

BCL10 functions in the CBM complex. Typically, CBM activation is mediated by CARD11 phosphorylation, inducing a confirmational change that recruits BCL10 to form signaling filaments that in turn activate scaffolding and caspase functions of MALT1 (49). *CARD11* is a well-described oncogene in ABC-DLBCL cases, acquiring activating mutations with potentially potent transforming effects (18,50–52). *BCL10* mutations have been only more recently recognized and minimally characterized by comparison. Xia and colleagues showed two classes of *BCL10* alteration occur, both activating in nature, serving either to stabilize or spontaneously activate formation of BCL10 filaments (6). We later showed these mutations drive resistance to multiple classes of targeted signaling inhibitor and that BTK inhibitor resistance could be overcome by co-treatment with venetoclax (5). The murine model system in this report, employing conditional expression of the activating BCL10 truncation S136X, now facilitates detailed further investigation. Among other findings is absent lymphoma onset in primary *Mb1.BCL10*^trunc^ animals, standing in marked contrast to expression of mutant *CARD11* from *Rosa26* at the same point in B-cell development. which resulted in fatal B-cell tumors within a week of birth (18). We therefore reveal significant heterogeneity in the effects of CBM mutational activation, dependent on the specific mechanism of activation.

Impaired GC formation was another intriguing finding we investigated in detail. By virtue of its association with the master GC regulator BCL6 in DLBCL pathogenesis, BCL10 GoF might be expected to potentiate GC reactions regardless of when it arises. Our data instead show that developmental timing not signaling strength determines the GC phenotype: Expression in NB cells (*Mb1.BCL10*^trunc^) promoted splenic follicles that failed to form GCs (**Fig. 3**), whereas the same allele restricted to cells already committed to the GC program (*C*_γ_*1.BCL10*^trunc^) produced expanded GCs (**Fig. 7**). However, when combined with *BCL6* overexpression, this timing dependence dissolved entirely for lymphomagenesis. In addition, after multiple stimulations, accumulated *BCL10*^trunc^-expressing B cells in *C*_γ_*1.BCL10*^trunc^ abrogated further GC responses. GC maintenance depends on BCL6 actively suppressing NF-kB/BCR signaling to preserve GCB cell viability, while CD40-NF-kB-IRF4 pathway reciprocally represses BCL6 transcription and promotes B cell exit from the GC reaction (53,54). Our transcriptional data recapitulated this circuit precisely: IRF4 expression was concomitantly upregulated with downregulated BCL6, consistent with premature GC exit. It’s worthwhile to note that overexpression of BCL6 in *Mb1.BCL10*^trunc^ couldn’t rescue the GC phenotype and therefore a failure of BCL6 function could not be a prime reason for the collapsed GC. We show through detailed single-cell analyses that *BCL10*^trunc^ expression in NB cells favors accumulation of pre-GC FOB cells over sustained GC participation (**Fig. 4**). Co-deregulation of BCL6, in addition to failing to rescue GC responses, drove accumulation of M3 MBCs, whose gene-expression signature was prominent in the lymphomas that developed in these animals and in corresponding patient tumors (**Fig. 6**). We therefore provide a second example of cooperating DLBCL oncogenes from the BN2/C1 group favoring transformation of non-GC MBCs, consistent with a relative absence of SHM signatures in these cases (21).

BCL10 GoF mutation in both our previous and current studies promote activation of specific gene-expression programs able to collaborate with BCL6 deregulation (**Fig. 6**). In particular, the phosphorylated STAT3 dimer binds directly to and up-regulates expression of BCL6 (42), and this is a consistent consequence of BCL10 mutational activation. We showed previously STAT3 activation occurs as part of a cytokine-mediated self-reinforcing signaling milieu initiated by BCL10 GoF (5). Here, we elaborate further on mediators of these effects with key consequences in the setting of BCL6 deregulation. First, STAT3 signaling is even further enhanced with the addition of BCL6 to BCL10 GoF. This leads in turn to further enhancement of *Bcl6* expression above that provided by the *Iµ-HABCL6* transgene alone without BCL10 GoF. These events occurred in parallel with prominent expression of IL9R, a marker also of the M3 MBC subset. We therefore illuminate the basis for frequent co-segregation of *BCL6* translocation with *BCL10* activating mutations in the pathogenesis of DLBCL. The findings reported herein not only elucidate the heterogeneity underlying DLBCL tumorigenesis but also reveal how a single gene mutation can redirect GC fate determination, even in the presence of its most well-established facilitator.

## Methods

### Mice and Cell Lines

All mouse studies were performed under the approval of the University of Miami Institutional Animal Care and Use committee. The mice were housed within a barrier facility. All equipment, food, or bedding exposed to the mice is autoclaved, irradiated (food), or chemically disinfected. All cages are handled under animal transfer stations (Biological Safety Cabinets). Mice are housed in individually ventilated, double sided, racks and are changed weekly. The mice were housed socially (2 or more mice per cage) with enrichment. For in vivo experiments, mice were selected based on their genotype, and all experiments were conducted using age-matched littermates. All experiments were conducted using mice aged 8 to 20 weeks except the survival curves or unless otherwise specified in the methods. For cell lines, RIVA (RRID:CVCL_1885) and TMD8 (RRID:CVCL_A442) cells were obtained from Deutsche Sammlung von Mikroorganismen und Zellkulturen (DSMZ) and cultured in RPMI (Gibco, 77975-356) with 10% FBS (35-010-CV, Corning), 1% penicillin-streptomycin (15140163, ThermoFisher). Cells were maintained in an incubator at 37*°*C with 5% CO_2_ and all cell lines were verified by STR finger printing and tested for mycoplasma.

### Conditional Knock In model

C57BL/6J (RRID:IMSR_JAX:000664) mice with an inducible human *BCL10*^S136X^ knock-in mutation on the *Rosa26* locus (Cyagen) were bred with *Mb1*-Cre mice (Jackson Laboratories) to generate a colony of *BCL10*^S136X^-*Mb1*-Cre (*Mb1.BCL10*^trunc^) mice, along with groups of heterozygous and homozygous mice for breeding and experimental controls. These mice were monitored regularly with weekly weights, monthly complete blood counts (CBCs) and flow cytometry and spleen volumes were measured by ultrasound every 3 months. An additional cohort was immunized every two weeks with chicken gamma globulin (003-000-002, Jackson ImmunoResearch) to stimulate an immune response. *Iµ-HABCL6* mice were a generous gift from Riccardo Dalla-Favera from the Institute for Cancer Genetics, Columbia University, New York. *Iµ-HABCL6* were bred with *Mb1.BCL10*^trunc^ to generate *BCL10*^S136X^-*Mb1*-Cre x *Iµ-HABCL6* (*BCL6.Mb1.BCL10*^trunc^) mice colony. *C*_γ_*1*-Cre mice were obtained from Dr. Martin Riva’s laboratory (University of Miami, Florida, USA) via an intramural transfer and crossed with *BCL10*^S136X^ knock-in mutation C57BL/6J mice to generate *BCL10*^S136X^-*C*_γ_*1*-Cre (*C*_γ_*1.BCL10*^trunc^) mice colony.

### PCR for Recombined Knock-In Allele

To check if the conditional knock in allele was recombined by Cre recombinase activity in the RSB-Mb1-cre mice, a PCR was performed on genomic DNA extracted from mouse tissue using the Qiagen AllPrep DNA/RNA extraction kit (80204). The PCR reactions were performed using Taq DNA polymerase. Primer sequences:

Forward 5’ – CCAGAATTGCGAGGTATCGCAT – 3’,

Reverse 5’ – ATAGATTCAACAAGGGTGTCCAG – 3’

### VDJ IgH PCR

B cells were harvested from the individual mice spleen through CD43 negative selection using Miltenyi columns as described earlier, and the cell pellets were washed two times with the cold PBS. Genome DNA was isolated using PureLink™ Genomic DNA Mini Kit (K182001, Thermo Fisher) as per manufacturer’s protocol. 50 ng of DNA was used as template and amplified using specific primers (55). PCR amplification was performed in 25 uL reaction mixture with the thermal cycling conditions consisting of an initial denaturation at 98 °C for 30 sec, followed by 33 cycles of denaturation at 98 °C for 10 sec, annealing at 67 °C for 30 sec, and extension at 72 °C for 90 sec. A final extension was performed at 72 °C for 10 minutes. Amplified product was resolved on one percentage agarose gel with 1X concentration of SYBR™ Safe DNA Gel Stain (S33102, Invitrogen) and pictures were captured at LI-COR imaging system (RRID:SCR_023227).

VHJ558-JH4 PCR (distal) (D)

Fw Primer VHJ558: gcgaagcttargcctgggrcttcagtgaag

Rev Primer JH4: aggctctgagatccctagacag

VHJ7183-JH4 PCR (proximal) (P)

Fw Primer VH7183: cggtaccaagaasamcctgtwcctgcaaatgasc

Rev Primer JH4: aggctctgagatccctagacag

Loading Control (L)

Fw Primer Cμ: tggccatgggctgcctagcccgggactt

Rev Primer Cμ: gcctgactgagctcacacaaggagga

### Quantitative PCR

The Maxima™ H Minus cDNA Synthesis Master Mix (ThermoFisher, M1661) was used according to the manufacturer’s protocols to synthesize cDNA from RNA extracted from mouse tissues using the Qiagen AllPrep DNA/RNA extraction kit (80204). qRT-PCR was performed using a StepOnePlus_TM_ Real-Time PCR System (ThermoFisher). Relative changes in expression were calculated using the comparative Ct (ΔΔCt) method and 18S rRNA was used as an internal control. All qPCR reactions were performed using PowerUp_TM_ SYBR. Green Master Mix (ThermoFisher, A25742).

Primer sequences:

<u>GFP #1</u>

Forward 5’ – AAGTTCATCTGCACCACCG – 3’

Reverse 5’-TCCTTGAAGAAGATGGTGCG – 3’

<u>GFP #2</u>

Forward 5’ – TGAGCAAAGACCCCAACGAG – 3’

Reverse 5’ – CTTGTACAGCTCGTCCATGC – 3’

### SRBC Immunization

SRBCs (Cocalico Biologicals, 20-1334A), were prepared for injections under aseptic conditions in a laminal flow hood. Briefly, 1 ml of citrated SRBCs was aspirated using an 18-gauze needle and transferred to a 50mL conical tube, which was then filled to volume (50 mL) with cold PBS. Cells were pelleted by centrifugation at 800g for 6 min at 4°C, and the supernatant was carefully discarded. This wash step was repeated once, or until the supernatant appeared relatively clear. The washed pellet was resuspended in cold PBS to a final volume of 4 mL and counted to confirm a final concentration of 1-2x 10^9^ cells/mL yielding a target immunization dose of 1-2x 10^8^ SRBCs in 100ul per animal. Route of administration was intraperitoneal. For repeated immunization, mice were immunized after every one month till they reached the endpoint.

### Generation of secondary recipient mouse models from genetically engineered mouse models

Lymphoma cells are isolated from enlarged sublingual lymph nodes of *Mb1.BCL10*^trunc^ mice and cultured on irradiated 3T3 feeder layer. Cells were periodically supplemented with the fresh B cell media. These cultured cells were transplanted into C57BL/6 (irradiated with sub-lethal dose) mice via intravenous injection and followed for the successful transplantation by tracking GFP in peripheral blood. Lymphoma cells grown in the recipients were extracted when abdominal distension was observed and were transplanted into new C57BL/6 recipients, as well as placed in culture *in vitro*. The lymphoma cell passaging in irradiated C57BL/6 mice was repeated; RPMI-1640 (Gibco, 77975-356), 15% fetal bovine serum FBS (35-010-CV, Corning), 2 mM L-glutamine (21051024, ThermoFisher), 10 mM HEPES (15630080, ThermoFisher), 1×penicillin-streptomycin (15140163, ThermoFisher), and 50μM 2-Mercaptoethanol (31350010, Thermo Fisher Scientific), as previously described.(56)

### B cell purification and ex vivo activation

8 to 12 weeks of mice were used for studying ex-vivo stimulation. Spleens were mashed and passed through 70-μm cell strainers (732-2758, VWR) to dissociate cells into single-cell suspensions. Red blood cells (RBCs) were lysed by resuspending in 2 ml lysis buffer (A1049201, Thermo Scientific) for 5 min at 25 °C. The lysis reaction was quenched by adding four volumes of B cell media (RPMI Medium 1640 1X HEPES without L-Glutamine (9159, FUJIFILM BIOSCIENCES INC) with 15% FBS (35-010-CV, Corning), 1% penicillin–streptomycin (15140163, ThermoFisher)), 1% L-Glutamine (21051024, ThermoFisher) and 55mM β mercaptoethanol (31350010, Thermo Fisher Scientific). Following RBC lysis, B cells were treated with DNase I (11284932001, Roche) for 15 minutes at RT and enumerated. Mature B cells were isolated by CD43 (Ly-48) MicroBeads (130-049-801, Miltenyi Biotec) in MACS buffer (500 ml PBS, 0.5% BSA, 0.5 M EDTA) followed by negative selection using LS Columns (130-042-401, Miltenyi Biotec). Before stimulation cells were labelled with Cell trace violet (CTV) (C34557, Thermo Fisher Scientific) for 20 minutes at 37° C in the dark. Excess CTV was removed by washing. Purified B cells were seeded at 1 × 10^6^ per ml of B cell media and cultured ex vivo with combinations of LPS (33.3μg ml^−1^; L4130, Sigma), IL-4 (12.5ng ml^−1^, 404-ML-010, R&D systems), F(ab′)_2_ anti-IgM (1µg ml^−1^;115-006-020, Jackson ImmunoResearch) and anti-CD40 monoclonal Antibody (HM40-3, Functional Grade, eBioscience) (1µg/mL; 16-0402-82, LIFE TECHNOLOGIES CORPORATION). Cells were analyzed at 48, 72 or 96 hours as indicated.

### Immunoblotting

For mouse samples B cells were isolated from the spleen using microbeads (CD43) through negative selection as mentioned above and selected B cells were washed and spun down at 800g for 5 min and the cell pellets were resuspended in Radioimmunoprecipitation assay (RIPA, C748H94, Thomas Scientific) buffer premixed with Halt™ Protease and Phosphatase Inhibitor Cocktail (78442, Thermo Fisher). The lysate was incubated at 95 °C for 5 min and then sonicated thrice for 10 s at 25% amplitude using a digital sonicator. After sonication, the lysate was spun at 15,000*g* for 5 mins and analyzed for protein concentration by Bradford Dye Reagent, Ready-to-Use solution (BioRad, 5000006). Lysates were run on gradient 4-12% SDS–PAGE gels and the resolved proteins were transferred at 26 V onto PVDF membrane (1620174, Biorad). The blots were blocked with 5% non-fat Blotting-Grade Blocker Milk (1706404, Bio-rad) and incubated with the primary and secondary antibodies as indicated. For the ABC cells lines, TMD8 and RIVA cells were induced with doxycycline (200ng/mL) 24 hours prior to collection of the cells, and the immunoblotting was performed as mentioned earlier (5). Antibodies: BCL6 (Cell signaling, 5650, RRID: AB_10949970, 1:1000), p-STAT3 (Cell signaling, 9145, RRID: AB_2491009, 1:500), STAT3 (Cell signaling, 4904, RRID: AB_331269, 1:1000), IL9r (Invitrogen, PA5-67619, RRID: AB_2688440, 1:500), anti-BCL10 (Cell signaling, 4237, RRID: AB_2228005, 1:1000) and Goat anti-rabbit HRP conjugated secondary antibody (ThermoFisher, 31460, RRID: AB_228341, 1:5000).

### Mouse embryonic fibroblasts (MEFs) isolation and BCl10 S136X expression

Embryos (Day 14.5) were dissected from the pregnant females from the RSB (human *BCL10*^S136X^ knock-in mutation on the *Rosa26* locus) and WT/WT female mice by removing the uterine horns under sterile conditions. The uterine horns were washed in DPBS (Fisher Scientific, 14190136) under aseptic conditions, followed by removing individual embryos from the uterine tissue. Before dissection of embryos, yolk sac and placenta were removed, and the exposed embryos were rinsed in sterile DPBS with antibiotic-antimycotic. Before tissue dissociation the embryos were decapitated, and fetal organs were removed. The individual embryos were kept in separate wells in a 6 well plate and tissues were minced. The minced tissues were incubated in trypsin overnight at 4°C in 15 ml falcon tube. The following morning, tubes were incubated at 37°C for 15 minutes to activate the trypsin and resuspended with 1 ml pipette and let it settle for 5 minutes. The undigested tissues were settled down, and the supernatant was transferred to a fresh tube. This supernatant was spun down at 1100 rpm for 5 min and pellet resuspend the cell pellet in 5 ml prewarmed MEF harvest medium T25 flask. Incubate for approximately 24 hours at 37°C, 5% CO2 under humid conditions. This was considered as passage 0 and the MEFs from passage 2 or maximum 3 were transduced with the virus generated using plasmid (MSCV-Cre, Addgene, RRID: 24064) expressing Cre recombinase engineered with mcherry using NEBuilder^®^ HiFi DNA Assembly Master Mix (New England Biolabs, E2621S). For generation of virus phoenix cells were transfected with MSCV-Cre-mcherry to generate the virus using Lipofectamine 3000 transfection reagent (ThermoFisher, L3000015) in Opti-MEM (Fisher Scientific, 11058-021). The medium was replaced by the DMEM (VWR, 45000-304) supplemented with 10 percent fetal bovine serum (FBS, Corning, 45000-736) after 12 hours of transfection). Supernatant with virus was harvested after 48 hours and 72 hours and passed through the 0.45-micron filters (VWR, 28143-312). 24 hours before transduction 1×10^5^ MEFs were plated in each well of 6 well plate. After 24 hours of plating and transducing the virus in DMEM supplemented with 2 percent of FBS and polybrene (4µg/ml, R&D systems, 77-111-0) without antibiotic. After 6 hours media was replaced with the DMEM supplemented with 10 percent FBS and 24 hours post transduction cells were observed for expression of mcherry from MSCV-Cre-mcherry plasmid and GFP from the RSB cassette. Pool of cells was harvested after 48 hours, and immunoblotting was performed as mentioned above (57). Antibodies; anti-mcherry (Antibodies.com, A85306, RRID:AB_2291286, 1:1000), anti-GFP (CST, 2956, RRID:1196615, 1:1000), anti-BCL10 (Cell signaling, 4237, RRID: AB_2228005, 1:1000), Goat anti-rabbit HRP conjugated secondary antibody (ThermoFisher, 31460, RRID: AB_228341, 1:5000).

### Tissue processing and single-cell preparation

Spleens and bone marrow were harvested from 12 weeks old male or female mice of each genotype at 7-day post SRBC immunization. Bone marrow was collected from femurs and tibias by flushing with FACS buffer. Spleens and bone marrow were dissociated into single-cell suspensions by mechanical disruption through 70µm cell strainer (Corning, 352350), and red blood cells were lysed in 1 ml of ACK lysis buffer (Thermo Fisher, A1049201) at for 5 minutes at room temperature. Lysis was quenched with 4 ml of FACS buffer and cells were spun down at 800g at 4°C. After centrifugation, supernatant was removed and the pellet was resuspended in 10 ml of FACS buffer and passed through 70µm cell strainer to remove any cell clumps and used for downstream analysis. Peyer’s patches (PPs) were mashed and passed through 70-µm cell strainer without RBC lysis and used for downstream analysis.

### Surface staining for flow cytometry

Cells were enumerated using trypan blue on automated counter (Invitrogen™ Countess™ 3 Automated Cell Counter) and 2×10^6^ cells per sample were washed twice in cold PBS. Cells were stained with either Zombie UV fixable viability dye (BioLegend, 423107, 1:1000) or Zombie UV yellow fixable viability dye (BioLegend, 423103, 1:800) for 15 minutes at room temperature, protected from light. The viability dye was quenched and washed out with FACS buffer, and cells were centrifuged at 800xg for 5 minutes at 4°C. Cells were then resuspended in FACS buffer containing anti-mouse CD16/32 Fc block (BD, 553142, 1:50) and incubated for 20 minutes at 4°C, protected from light. Following Fc blocking, cells were spun down and supernatant was removed and cells were resuspended in the 100µl of antibody cocktail. After staining, cells were washed twice in FACS buffer to remove unbound antibodies and resuspended in 200µl of FACS buffer. Surface staining was performed with antibodies against CD19, CD3, Cd11b, Gr-1, CD93, CD19, CD21, CD138, CD23, CD24, CD43, BP-1, B220, FAS, GL7, IgD, anti-IgM, IgG1; clones’, fluorophores, vendors, catalog numbers, and RRIDs listed in Supplementary Table 1.

### CD8 T cell exhaustion markers

Unimmunized mice aged 4-6 months were euthanized and spleens were harvested. Single-cell suspensions were prepared, and viable cells were enumerated using trypan blue. T cells were enriched by negative selection using EasySep^Tm^ mouse T cell isolation kit (STEMCELL Technologies, 19851) according to manufacturer’s instructions. Enriched cells (1×10^6^ per sample) were stained for flow cytometry as described above, using the antibodies listed in Supplementary Table 1.

### Apoptosis assay

96 hours after anti-IgM stimulation, cells were harvested, washed twice in cold PBS, and resuspended in 100µl of 1X Annexin V binding buffer (BD, component number: 51-66121E). Cells were stained with Annexin-V (Biolegend, 640949, 5µl/100ul reaction) and propidium iodide (BD, component number:51-66211E) for 15 minutes at room temperature in the dark, then analyzed immediately without fixation.

### Flow data acquisition and analysis

Spleen, lungs and bone marrow cells were acquired on spectral Cytek Aurora or BD FACSymphony A5SE. Peyer’s’ patches analysis was done on BD FACSCanto II. Ex-vivo B cell stimulation and engrafted mice tumor cells’ flow data was acquired on Cytek Aurora spectral cytometer. For spectral unmixing single stain reference controls were prepared using either the cells or UltraComp eBeads (Thermo Scientific, 01-2222-42). Unstained cells were used for the extraction of autofluorescence. For the EGFP control mutant B cells were used as reference control throughout flow experiments. For conventional flow cytometer single stained cells were used as reference controls along with the unstained cells from WT/WT or the *Iµ-HABCL6* genotype. Absolute cell numbers were counted by adding 10µl of CountBright absolute counting beads (Invitrogen, C36950) to the 200 µl of cell volume for counting of absolute number of the cells. Data was analyzed using FlowJo (BD, Version 10.10.0, RRID: SCR_008520).

### Immunohistology

Tissue samples (spleen, lungs and tibia) were fixed in 10% neutral-buffered formalin (VWR, 89370-094), for 48 hours at room temperature and processed in alcohol and xylene, and fixed tissues were embedded in paraffin wax. For histopathological analysis, specimens were evaluated by a board-certified veterinary pathologist and hemopathologist. Sample preparation, processing, and H&E staining were performed as previously described. All the staining was performed, using the BOND RX protocol (Leica Microsystems, RRID:SCR_025548). In brief the FFPE sections were pre-baked at 60 °C for 30 minutes and loaded at automated Stainer Leica Bond Rx to perform the standard bond polymer refine IHC protocol which included dewaxing and rehydration. Antigen retrieval was done with citrate buffer pH 6.0 (Bond ER1 solution for 20 minutes followed by peroxidase block for 15 minutes. The slides were further blocked in 10 % normal goat serum for 30 minutes. After blocking slides were incubated with the primary antibody for 30 minutes. diaminobenzidine (DAB) staining was performed with Bond Mixed Refine for 10 minutes and counterstained with light hematoxylin for 5 minutes. The excess stain was washed, and slides were automated dehydrated at Leica ST5020 and automated coverslip at Leica CV5030 and observed under microscope. All the histology image quantifications were done using ImageJ software. All imaging was done at RT using a Leica microsystem (CMS GmbH) or the EVOS inverted microscope (Invitrogen). Antibodies used for histology: BCL6 (NovusBiologicals, NBP3-07526, 1:1000), CD20 (Cell signaling, 70168, 1:400, RRID:AB_2943602), CD10/Neprilysin (Cell signaling, 58145, 1:300), IRF4 (Cell signaling, 62834, 1:600, AB_2877647), B220 (NovusBiologicals, NBP3-11980, 1:500), CD19 (Cell signaling, 90176S, 1:600, RRID:AB_2800152), GFP (Abcam, ab183734, 1:200), Ki67 (Cell signaling, 12202S, 1:500, RRID:AB_2620142) and CD3 (Cell signaling, 99940, 1:300, AB_2755035).

### Cell Preparation and GEM-X Flex Fixation

Single cell splenocytes were prepared by carefully extracting spleens from (WT, *Mb1.BCL10*^trunc^, *C*_γ_*1.BCL10*^trunc^, *Iµ-HABCL6* and TH) as explained above. Dead cells were depleted using the EasySep™ Annexin V-based Dead Cell Removal Kit (STEMCELL Technologies, 17899), following standard manufacturer instructions. High-quality single cell splenocytes (90% viable cell) were fixed using 10X Genomics GEM-X Flex sample Preparation v2 Kit (10X Genomics, PN-1000781) according to the manufacturer’s direction. Briefly, the splenocytes were resuspended in 1 ml of cold 0.04% BSA (Tribioscience, TBS8031) in PBS and centrifuged at 300 × g for 5 min at 40 C. The washed cells were resuspended with 0.5 ml Fixation Buffer-B followed by overnight (16-24 h) incubation at 40 C. After the incubation, 0.5 ml of Additive-C was added to the cells in Fixation buffer-B and mixed the reaction by pipetting 5 times followed by centrifugation at 850 × g for 5 min at room temperature. The supernatants were removed without disturbing the pellet and resuspended them in 1 ml chilled Quenching Buffer-B by pipetting 5 times and maintained on ice. The cell concentration and quality of the fixed samples was measured using Nexcelom Cellometer K2 (Nexcelom Bioscience) by employing ViaStain AOPI Staining Solution (Revvity). After confirming the quality and concentration of the fixed cells, 100 µL of pre-warmed enhancer (at a 1:10 volume ratio) and 275 µL of 50% glycerol (to achieve a final glycerol concentration of 10%) were added to each sample. The samples prepared were then stored at −80°C until further processing using the GEM-X Flex Gene Expression protocol.

### Quality control, cell-type clustering, and major cell-type identification

ScRNA-sequencing data were processed using Cell Ranger (v7.0.0, 10x Genomics) to generate gene expression matrices aligned to the mm10 reference genome. Quality control and downstream analysis were performed using Seurat (v5.0.0) in R (v4.3.0), filtering cells with <200 or >6000 expressed genes, <500 UMIs, or >20% mitochondrial reads, and removing doublets identified by scDblFinder. Data were normalized using SCTransform with regularization, and batch effects between samples were corrected using Harmony on the first 30 principal components. Graph-based clustering was performed using the Louvain algorithm (resolution=0.6) on a shared nearest neighbor graph (k=20), with visualization via UMAP. Cell types were annotated using SingleR with the Immgen reference database, validated by expression of canonical markers. For B cell subset analysis, an additional clustering step was performed at resolution 1.2, which identified 28 distinct B cell clusters that were annotated based on marker gene expression patterns characteristic of B cell developmental stages (NB, GC, LZ, DZ, memory, and plasma cells). RNA velocity was estimated using scVelo to infer transcriptional dynamics and predict future cell states by modeling splicing kinetics (58). Differential expression analysis between genotypes within each cell type was performed using limma and edgeR incorporating cellular detection rate as a covariate (adjusted p < 0.05, |log2FC| > 0.5) with pseudo-bulk differential expression via Libra. Functional pathway enrichment was evaluated by GSEA using Hallmark gene sets from MSigDB conducted to identify functional pathways altered between genotypes. Cell-cell communication networks were inferred using CellChatv2 following the published protocol (36), with the underlying inference model as described previously (59).

### Data and statistical analysis

Results are presented as mean□±□s.e.m., mean ± s.d., and mean only wherever applicable. *P* values were calculated by unpaired, two-tailed *t*-test or by two-way ANOVA with \**P*□≤□0.05, \*\**P*□≤□0.01, \*\*\**P*□≤□0.001, \*\*\*\**P*□≤□0.0001 unless otherwise specified in the methods or legends. Analysis was done by using GraphPad software version 10.6.1(892), RRID:SCR_002798. Experimental and genotyping schematics were generated using BioRender (licensed).

## Data Availability

All data in this study were generated by the authors or were analyzed from primary sources cited and are available as needed upon request from the corresponding author. Single-cell RNA sequencing data are publicly available at the Gene Expression Omnibus (https://www.ncbi.nlm.nih.gov/geo/) under accession #GSE334941.

## Authors’ contributions

Contribution: D.C. performed all the bench and mice experiments. AAY analyzed single cell RNA-seq data under the supervision of F.M. and J.H.S. V.K., and J.C. provided input in analyzing and designing ex-vivo stim experiments. H.T.H. and D.B. maintained the mice colony and did SRBC injections. C.A. interpreted in vivo pathology. C.A.C., V.S, M.L., and D.H. assisted in mice necropsy and single cell sample preparation. D.C. and J.H.S. wrote and edited the manuscript with input from all authors.

## Acknowledgements

The authors gratefully acknowledge Mr. Steve Dwoskin’s generous support of this project. Research in this publication was performed in part at the Cancer Modeling and Onco-Genomic Shared Resources (CMSR RRID: SCR022891, OGSR RRID: SCR022502) of the Sylvester Comprehensive Cancer Center at the University of Miami, which is funded by the USA National Institutes of Health, National Cancer Center under P30CA240139.

