## Supplementary data for "*BCL10* Gain-of-Function Impairs Germinal Center Participation by Naïve B Cells and Cooperates with *BCL6* to Drive Extrafollicular Diffuse Large B-Cell Lymphoma"

### Supplementary Appendix

#### Table of Contents


***SUPPLEMENTARY TABLES***
**Table S1.** Antibody List

***SUPPLEMENTARY FIGURES***
**Supplemental Figure 1**. Characterization of *Mb1.BCL10*$^{trunc}$ mice.
**Supplemental Figure 2**. BCL6 Synergizes with BCL10 GoF in Lymphomagenesis.
**Supplemental Figure 3**. *BCL10*$^{S136X}$ imposes a block in germinal center formation not overcome by BCL6.
**Supplemental Figure 4**. *BCL10*$^{trunc}$ Promotes Co-Stimulation-Independent Proliferation.
**Supplemental Figure 5**. Single-cell atlas of splenic immune populations from SRBC immunized mice across five genotypes**.**
**Supplemental Figure 6.** Single cell landscape of CD4 positive and CD8 positive population across genotypes.
**Supplemental Figure 7.** Transcriptional heterogeneity and genotype-dependent composition of splenic memory B cell in SRBC immunized mice.
**Supplemental Figure 8.** Repetitive SRBC immunization of *Cy1.BCL10*$^{trunc}$ lead to abrogated GCs.


***SUPPLEMENTARY TABLES***

**Table S1.** Antibody List

| Serial No. | Name of the Antibody | Catalog | Fluorophore | Clone | RRIDs | Provider | Application |
|---|---|---|---|---|---|---|---|
| 1 | CD3 | 100218 | PerCP/Cy5.5 | 17A2 | AB_1595492 | Biolegend | Flow |
| 2 | CD11b | 101228 | PerCP/Cy5.5 | M1/70 | AB_893232 | Biolegend | Flow |
| 3 | Gr-1 | 108428 | PerCP/Cy5.5 | RB6-8C5 | AB_893558 | Biolegend | Flow |
| 4 | CD93 | 12-5892-82 | PE | AA4.1 | AB_466018 | Invitrogen | Flow |
| 5 | CD19 | 115554 | PE/Dazzle 594 | 6D5 | AB_2564001 | BioLegend | Flow |
| 6 | CD21 | 123420 | PE/Cy7 | 7E9 | AB_1953277 | BioLegend | Flow |
| 7 | CD23 | 101642 | APC/Fire 750 | B3B4 | AB_2904272 | BioLegend | Flow |
| 8 | IgD | 744291 | BV421 | 11-26c.2a | AB_2742121 | BD | Flow |
| 9 | IgM | 743326 | BV650 | II/41 | AB_2741427 | BD | Flow |
| 10 | B220 | 563894 | BV786 | RA3-6B2 | AB_2738472 | BD | Flow |
| 11 | CD24 | 101838 | PE/Dazzle594 | M1/69 | AB_2566732 | BioLegend | Flow |

| | | | | | | | |
|---|---|---|---|---|---|---|---|
| 12 | BP-1 | 17-5891-82 | APC | 6C3 | AB_2762698 | Invitrogen | Flow |
| 13 | CD19 | 557399 | APC-H7 | 1D3 | AB_1645234 | BD | Flow |
| 14 | IgM | 743329 | BUV395 | II/41 | AB_2741430 | BD | Flow |
| 15 | IgG1 | 748403 | BUV737 | X56 | AB_2872822 | BD | Flow |
| 16 | IgD | 405731 | Alexa Fluor 700 | 11-26c.2a | AB_2563341 | Biolegend | Flow |
| 17 | 7AAD | 420404 | | | | Biolegend | Flow |
| 18 | CD8 | 612898 | BUV805 | 53-6.7 | AB_2870186 | BD | Flow |
| 19 | CD4 | 569182 | BUV563 | RM4-5 | AB_3684848 | BD | Flow |
| 20 | TIGIT | 142111 | BV421 | 1G9 | AB_2687311 | Biolegend | Flow |
| 21 | PD-1 | 109109 | PE/Cy7 | RMP1-30 | AB_572016 | Biolegend | Flow |
| 22 | IgG3 | 749009 | BUV563 | R40-82 | Not confirmed | BD | Flow |
| 23 | IgA | 1040-09 | PE | | AB_2794375 | SouthernBiotech | |

#### Supplemental Figures with Legends:

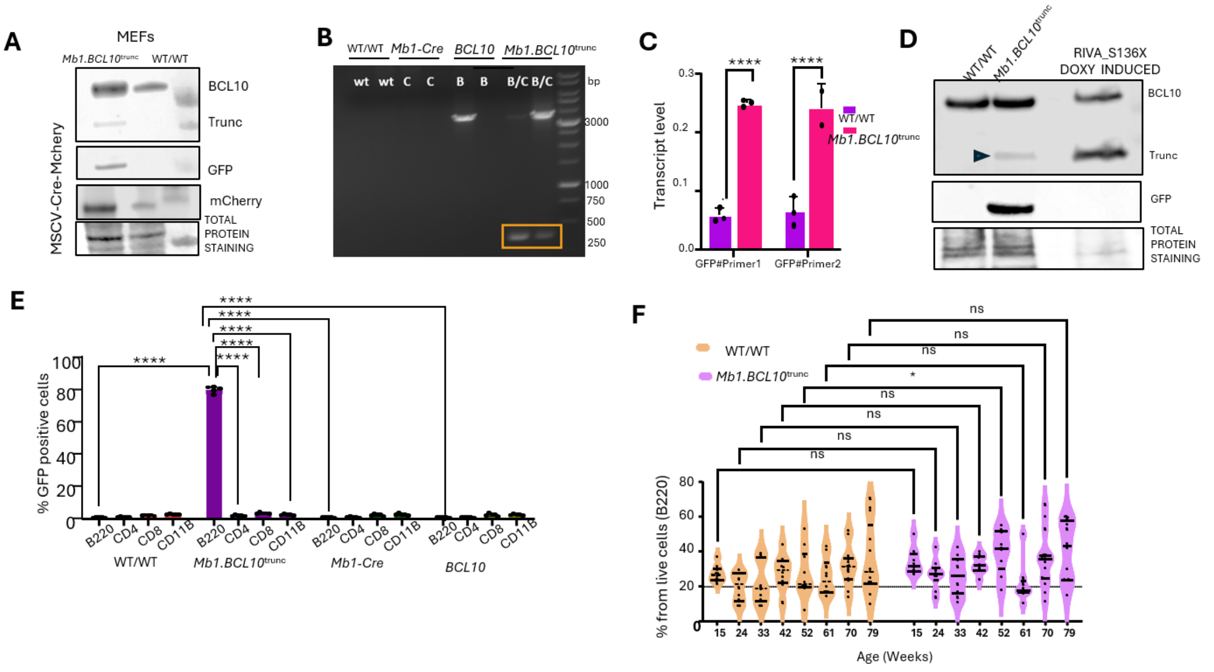

G

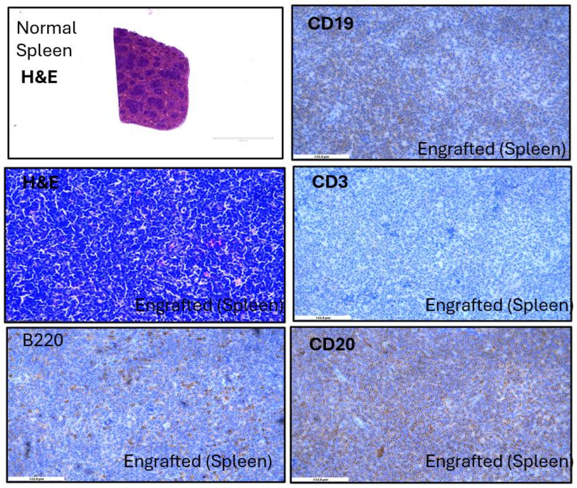


H

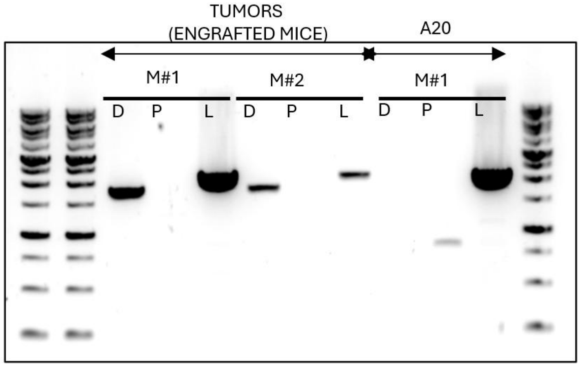


**Supplementary Figure 1. Characterization of *Mb1.BCL10*$^{trunc}$ mice; A.** Western blot showing truncated BCl10-S136X protein, GFP, mcherry in MEFs isolated from WT and *Mb1.BCL10*$^{trunc}$ embryos. **B.** PCR results confirming for recombined allele which is approximately 360bp. **C.** Quantitative PCR results of DNA from mice of different genotypes. GFP is only expressed when the mutant is present and therefore the presence of GFP indicates expression of the mutation **D**. Western blot showing truncated BCL10 protein (Lower band and marked with arrow) in the pure B cell population isolated from the spleen of WT or *Mb1.BCL10*$^{trunc}$. RIVA cells with doxycycline inducible overexpression of truncated BCL10 were used as a positive control. **E**. Flow cytometry analysis of peripheral blood of mice of all four genotypes, looking specifically at B220 positive cells among different genotypes. **F**. Flow cytometric analysis for B220, CD4, CD8 and CD11b of peripheral blood during the different time courses (12- 80 weeks).**G**. H&E for spleen for control (WT/WT) and H&E and IHC for the spleen isolated from engrafted mice. IHC markers: CD3, B220, CD20, CD19. **H**. VDJ PCR clonality showing the monoclonal population (single band) of the tumor cells isolated from the spleen of transplanted mice. A20 mouse tumor cell line was used as a positive control. D stands for distal region and P stand for proximal region amplifying diversity in $V_H$ family of genes. L stands for loading control amplifying the Cµ region. M stand for the mouse numbers.

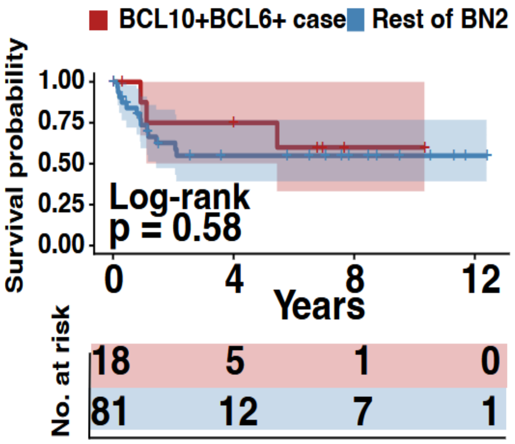


**Supplementary Figure 2. BCL6 Synergizes with BCL10 GoF in Lymphomagenesis;** Survival probability, all 18 human patients' cases harboring *BCL10+BCL6+* translocation/mutations vs rest of cohort in BN2 group (n=81). Log-rank p = 0.58 (vs BN2 cases only) using Lymphgen dataset.

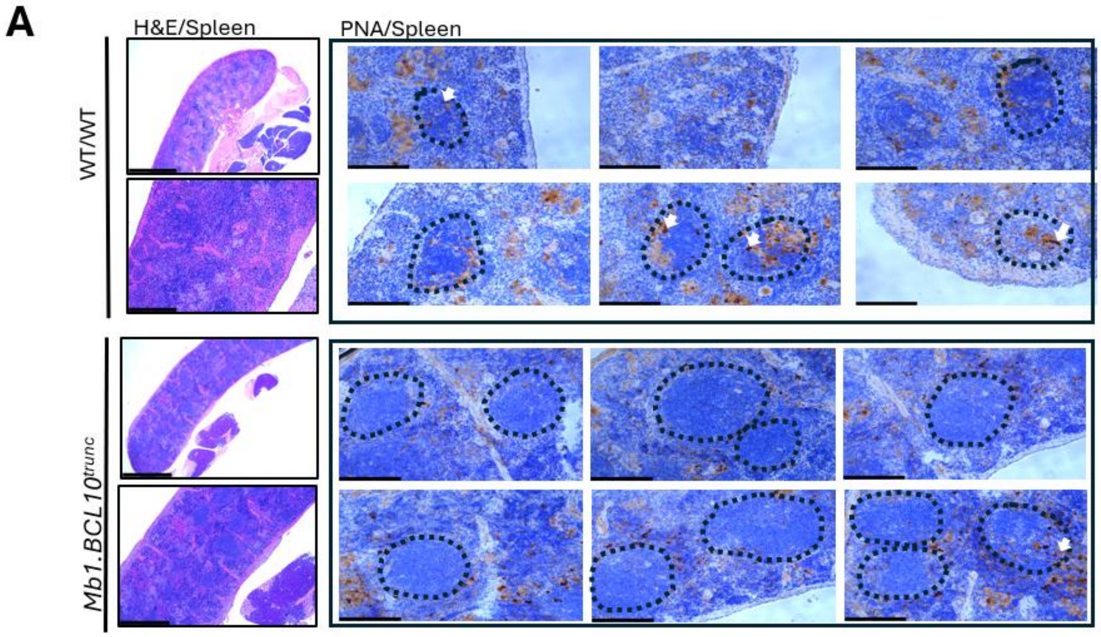

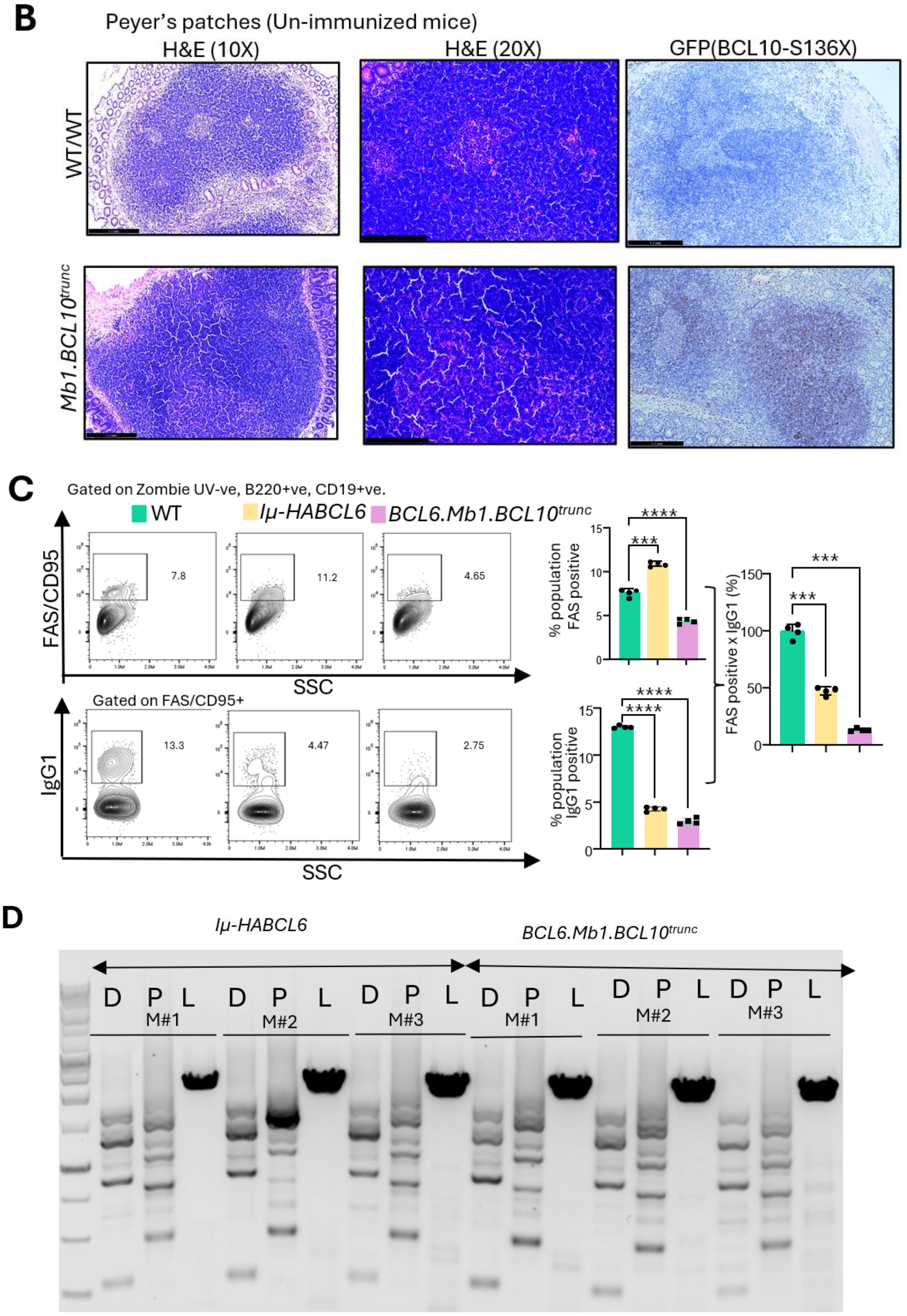
B
Peyer's patches (Un-immunized mice)
H&E (10X)
H&E (20X)
GFP(BCL10-S136X)
WT/WT
Mb1.BCL10trunc
C
Gated on Zombie UV-ve, B220+ve, CD19+ve.
WT
Iμ-HABCL6
BCL6.Mb1.BCL10trunc
FAS/CD95
SSC
7.8
11.2
4.65
Gated on FAS/CD95+
IgG1
13.3
4.47
2.75
% population FAS positive
% population IgG1 positive
FAS positive x IgG1 (%)
D
Iμ-HABCL6
BCL6.Mb1.BCL10trunc
D P L
M#1
M#2
M#3

**Supplementary Figure 3. *BCL10*$^{S136X}$ imposes a block in germinal center formation not overcome by BCL6; A.** H&E and PNA staining for the spleens isolated from mice immunized with CGG in WT/WT and *Mb1.BCL10*$^{trunc}$. Black dotted lines indicate the follicles. White arrow indicates the PNA staining. **B.** H&E and GFP (correspond to truncated BCL10$^{S136X}$) for Peyer's patches of WT/WT and *Mb1.BCL10*$^{trunc}$ in unimmunized mice. GFP annotate truncated BCL10-S136X. **C.** Representative FACS plots (left panel) and the quantification plots (right panel) showing the GL7 and FAS positive population across different genotypes (WT/WT, *Iμ-HABCL6* and *BCL6.Mb1.BCL10*$^{trunc}$) isolated from SRBC immunized mice and IgG1was used to quantify class switched cells among GL7 and FAS positive population. **D.** VDJ PCR clonality assay for the B cells isolated from the spleen of *BCL6.Mb1.BCL10*$^{trunc}$ and *Iμ-HABCL6*. M represents the different mice numbers. D stands for distal; P stands for proximal ends of the VDJ chain and L stands for loading control amplifying the Cμ region.

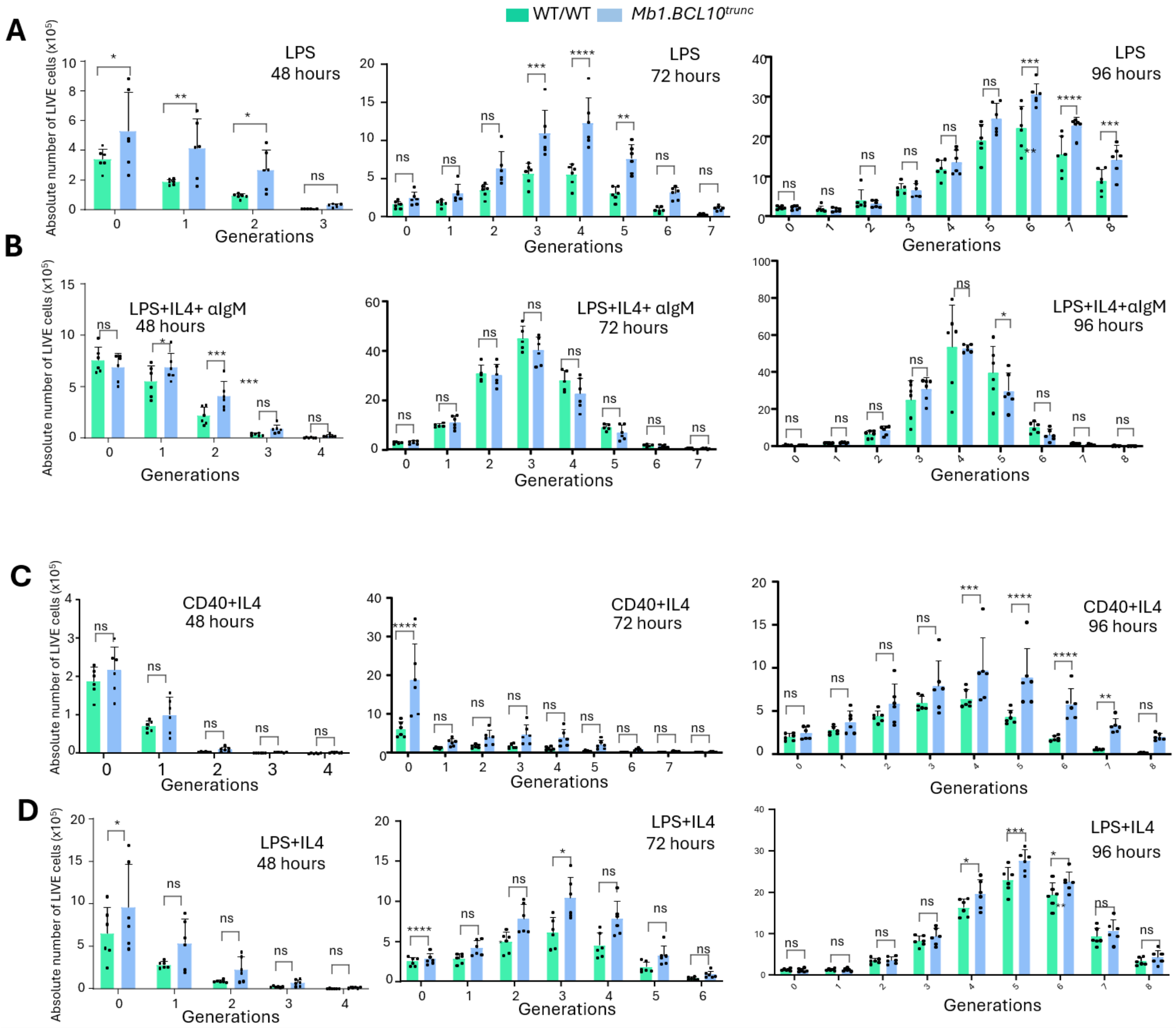

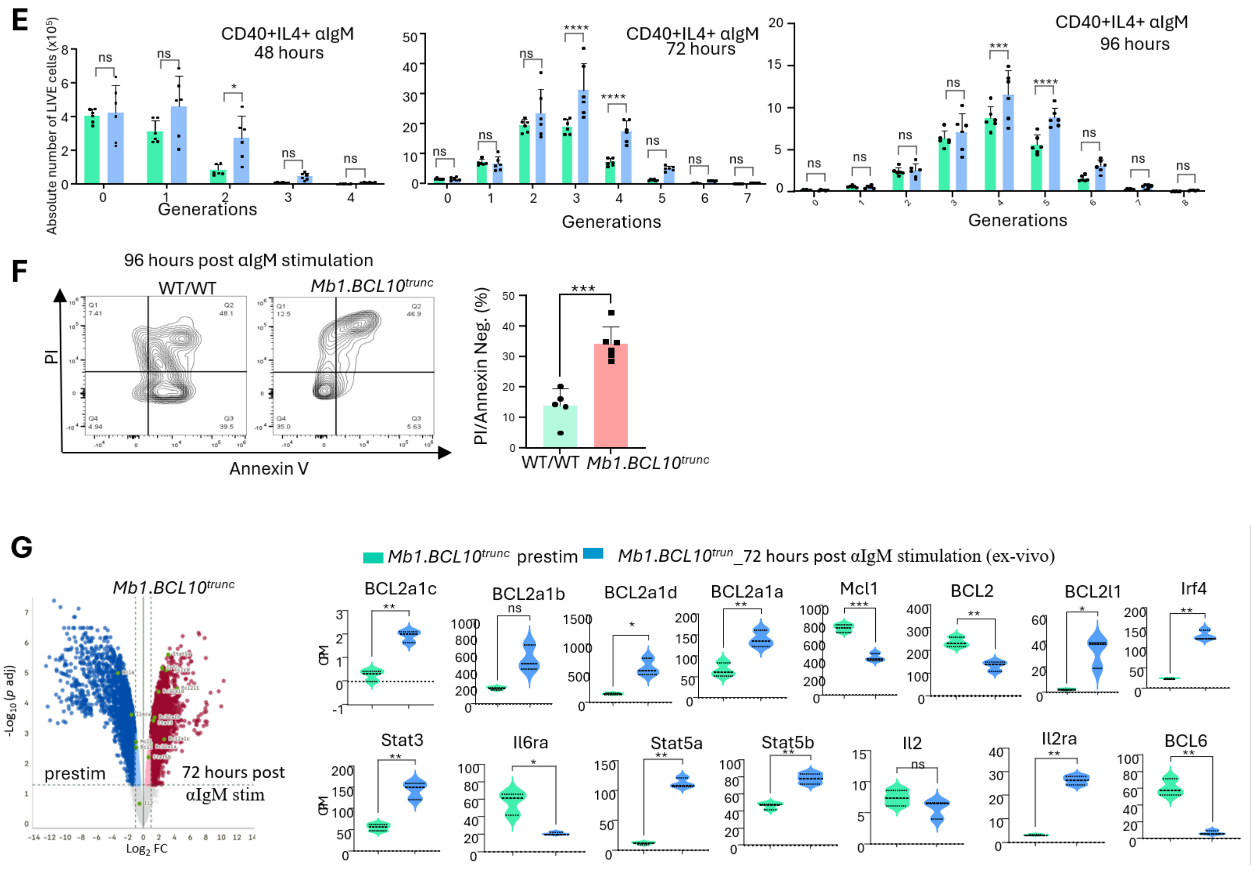


**Supplementary Figure 4. *BCL10*$^{\textbf{trunc}}$ Promotes Co-Stimulation-Independent Proliferation; A**. Bar graphs show the absolute count of B cells for each generation at 48, 72 and 96 hours post stimulation with different stimulants in ex-vivo culture; **A.** LPS. **B.** LPS+IL4+αIgM. **C**. CD40+IL4 **D**. LPS+IL4 **E**. CD40+IL4+αIgM. **F**. Representative FACS plots from apoptosis assay and bar plots showing the percentage of B cells negative from PI and annexin V post 96 hours of αIgM stimulation in ex-vivo culture. **G**. Volcano plot from bulk RNA sequencing from the cells harvested 72 hours of αIgM stimulation in ex-vivo culture in *Mb1.BCL10*$^{trunc}$. Cells before stimulation (prestim) were taken as control for the comparison. Bar plots showing the genes deregulated after αIgM stimulation in *Mb1.BCL10*$^{trunc}$ . Unpaired t test was used to calculate statistical significance $*P \leq 0.05$, $**P \leq 0.01$, $***P \leq 0.001$.

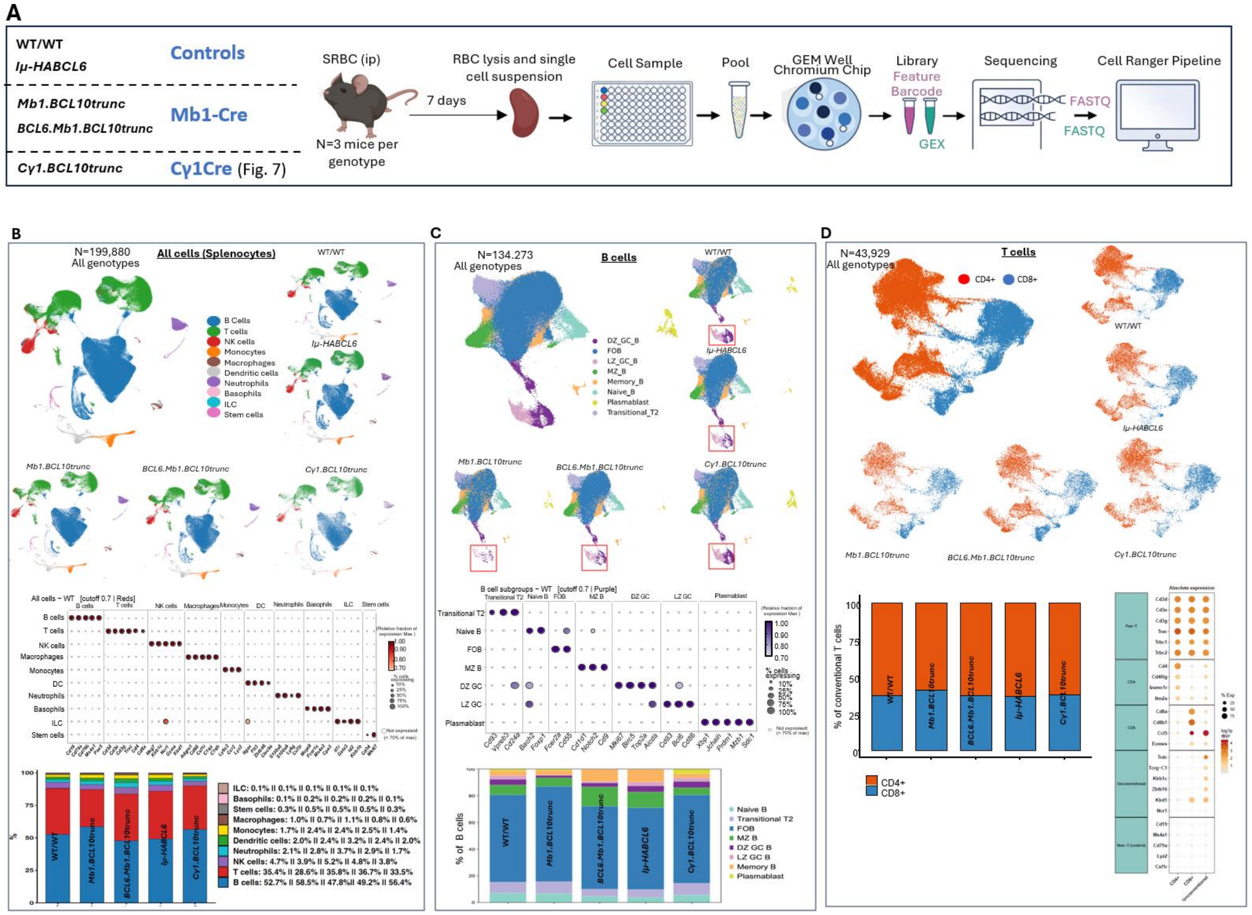


**Supplementary Figure 5. Single-cell atlas of splenic immune populations from SRBC immunized mice across five genotypes.** Experimental design. WT/WT, *Mb1.BCL10*$^{\text{trunc}}$, *BCL6.Mb1.BCL10*$^{\text{trunc}}$ , *Iμ-HABCL6* and *Cγ1.BCL10*$^{\text{trunc}}$ mice (n=3 per genotype). **B**. UMAP projection of 199,880 sequenced cells (splenocytes), colored by the ten major populations identified (B cells, basophils, dendritic cells, ILC, macrophages, monocytes, neutrophils, NK cells, stem cells and T cells). Dot plots of canonical marker genes used to annotate the major populations. Stacked bar plot of major immune-cell proportions per genotype. **C.** UMAP of the re-clustered 134,273 B cells colored by eight major clusters (Naïve, FOB, LZ, DZ, MZ, Memory, Transitional and Plasmablast B cells) compartment for each genotype, (WT/WT, *Mb1.BCL10*$^{\text{trunc}}$, *BCL6.Mb1.BCL10*$^{\text{trunc}}$ , *Iμ-HABCL6* and *Cγ1.BCL10*$^{\text{trunc}}$). Dot plots of canonical markers (*CD93, CD24a, Bach2, Foxp1, Fcer2a, CD55, Cd1d1, Notch2, CD9, Mki67, Birc5, Top2a, Aicda, CD83, Bcl6, CD86, Xbp1, J chain, Prdm1, Mzb1* and *sdc1*) used to annotate the clusters of B cell populations. Stacked bar plot of major B cell proportions per genotype. Red boxes marked the GC population decreased in *Mb1.BCL10*$^{\text{trunc}}$ and *BCL6.Mb1.BCL10*$^{\text{trunc}}$. **D**. UMAPs of the re-clustered T-cell compartment using 43,929 T cells grouped and individually for each genotype, (WT/WT, *Mb1.BCL10*$^{\text{trunc}}$, *BCL6.Mb1.BCL10*$^{\text{trunc}}$, *Iμ-HABCL6* and *Cγ1.BCL10*$^{\text{trunc}}$). Dot plots of canonical markers are used to annotate the CD4 positive and CD8 positive clusters of T cell populations. Stacked bar plot showing proportions of CD4 positive and CD8 positive T cell population per genotype.

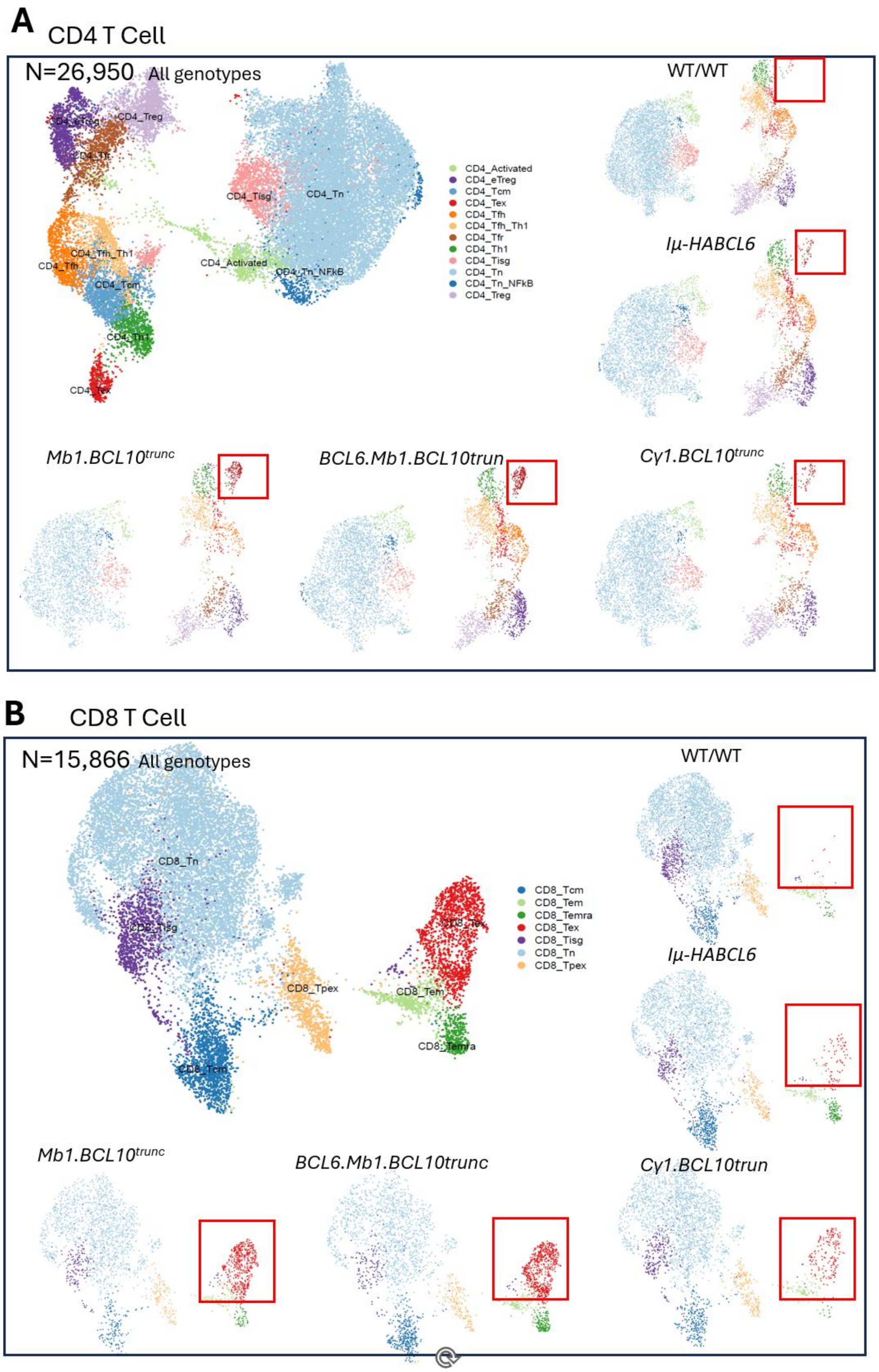
A
CD4 T Cell
N=26,950 All genotypes
CD4_Treg
CD4_Tisg
CD4_Tn
CD4_Tfh_Th1
CD4_Tfh
CD4_Activated
CD4_Tn_NFkB
CD4_Tcm
CD4_Th1
CD4_Tex
CD4_Activated
CD4_eTreg
CD4_Tcm
CD4_Tex
CD4_Tfh
CD4_Tfh_Th1
CD4_Tfr
CD4_Th1
CD4_Tisg
CD4_Tn
CD4_Tn_NFkB
CD4_Treg
WT/WT
Iμ-HABCL6
Mb1.BCL10trunc
BCL6.Mb1.BCL10trun
Cγ1.BCL10trunc
B
CD8 T Cell
N=15,866 All genotypes
CD8_Tn
CD8_Tisg
CD8_Tpex
CD8_Tem
CD8_Temra
CD8_Tcm
CD8_Tcm
CD8_Tem
CD8_Temra
CD8_Tex
CD8_Tisg
CD8_Tn
CD8_Tpex
WT/WT
Iμ-HABCL6
Mb1.BCL10trunc
BCL6.Mb1.BCL10trunc
Cγ1.BCL10trun

C

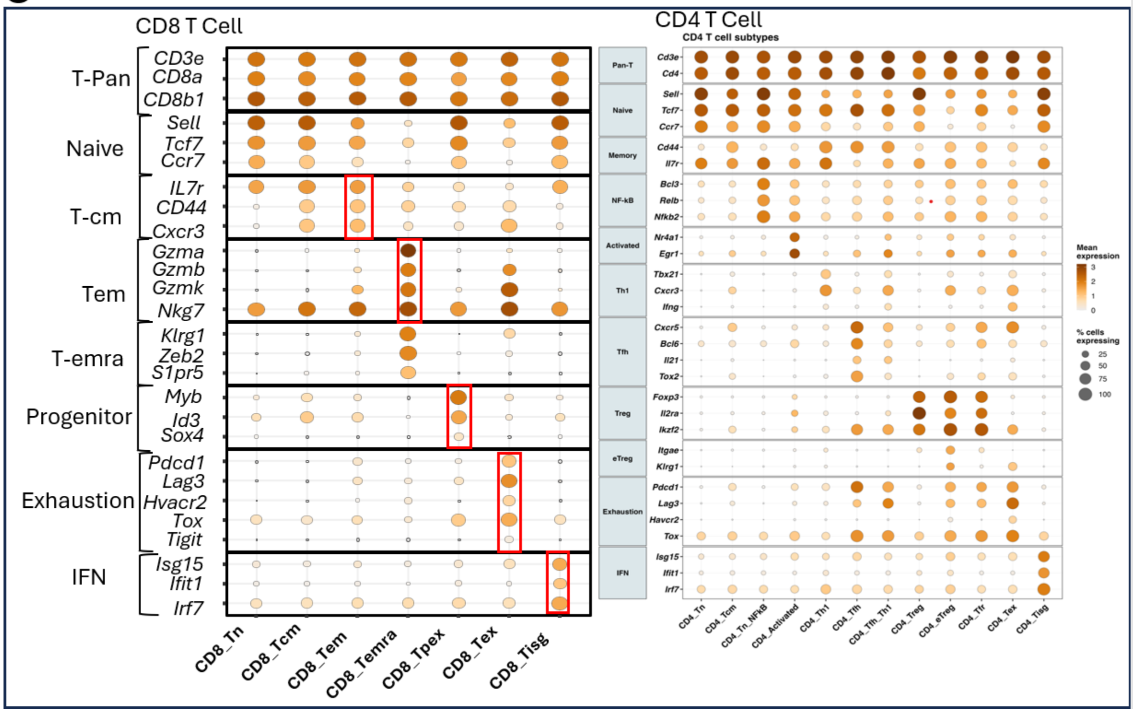


D

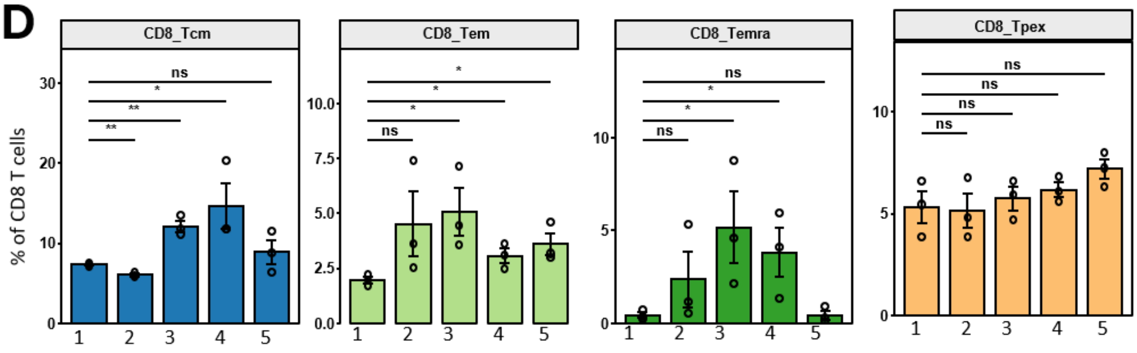


**1.** WT/WT **2.** *Mb1.BCL10trunc* **3.** *BCL6.Mb1.BCL10trunc* **4.** *Iμ-HABCL6* **5.** *Cγ1.BCL10trunc*

**E** CD4 Subtypes

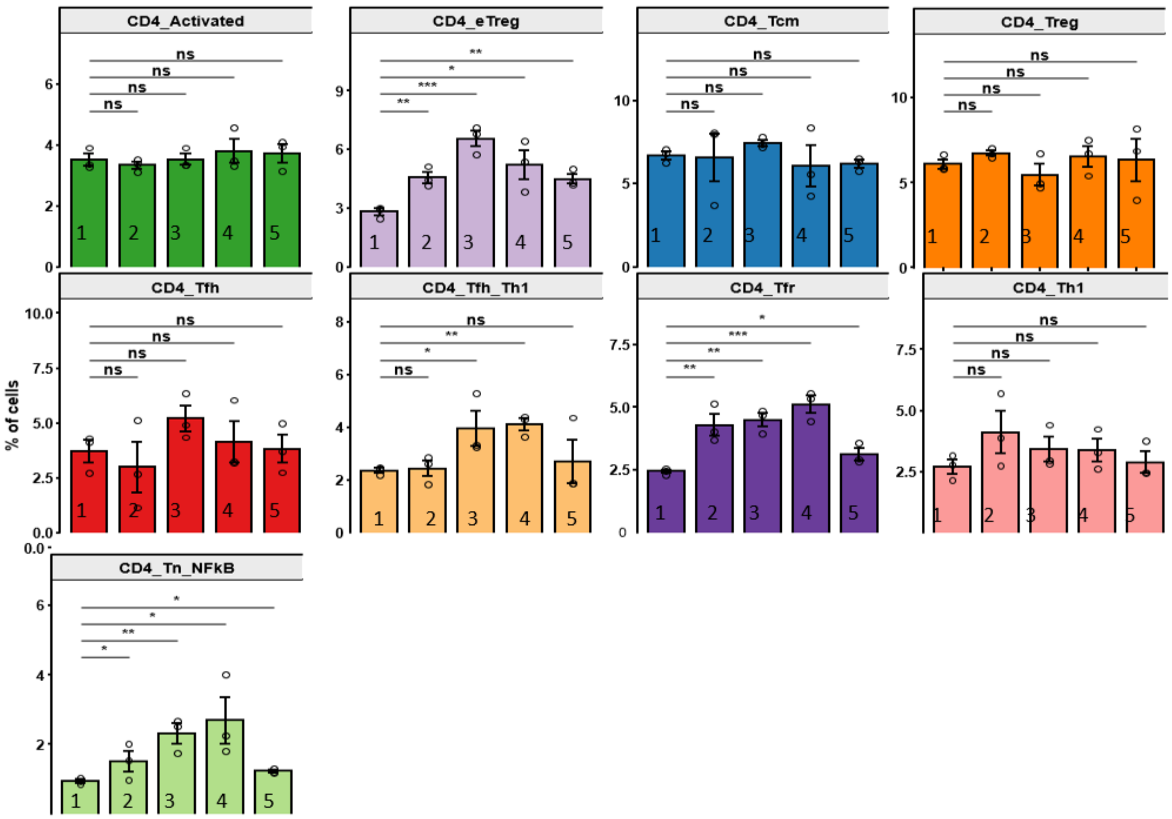


**1.** WT/WT **2.** *Mb1.BCL10trunc* **3.** *BCL6.Mb1.BCL10trunc* **4.** *Iμ-HABCL6* **5.** *Cγ1.BCL10trunc*

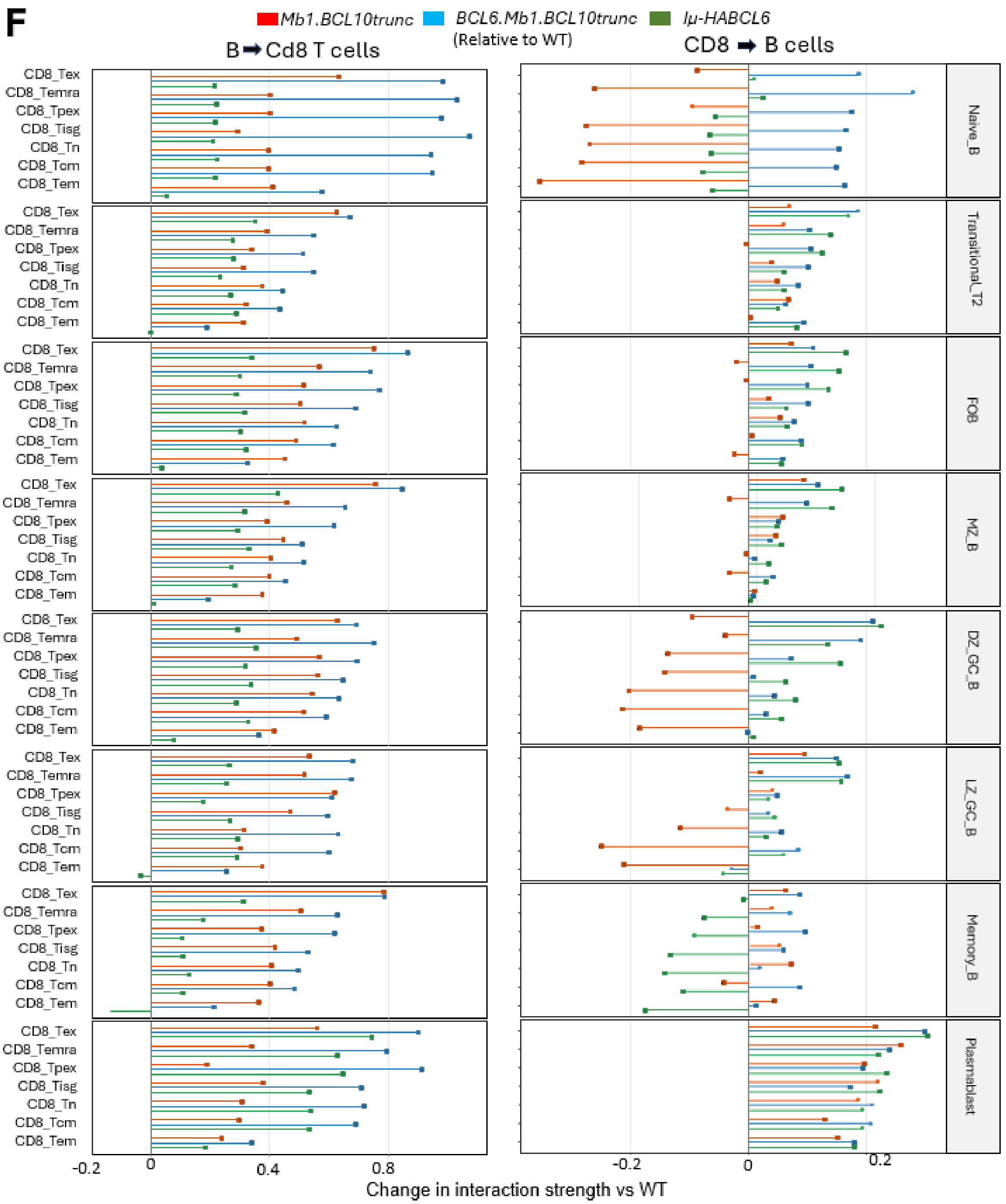
F
Mb1.BCL10trunc
BCL6.Mb1.BCL10trunc
Iμ-HABCL6
(Relative to WT)
B ➡ Cd8 T cells
CD8 ➡ B cells
CD8_Tex
CD8_Temra
CD8_Tpex
CD8_Tisg
CD8_Tn
CD8_Tcm
CD8_Tem
Naive_B
Transitional_T2
FOB
MZ_B
DZ_GC_B
LZ_GC_B
Memory_B
Plasmablast
-0.2
0
0.4
0.8
-0.2
0
0.2
Change in interaction strength vs WT

**G**

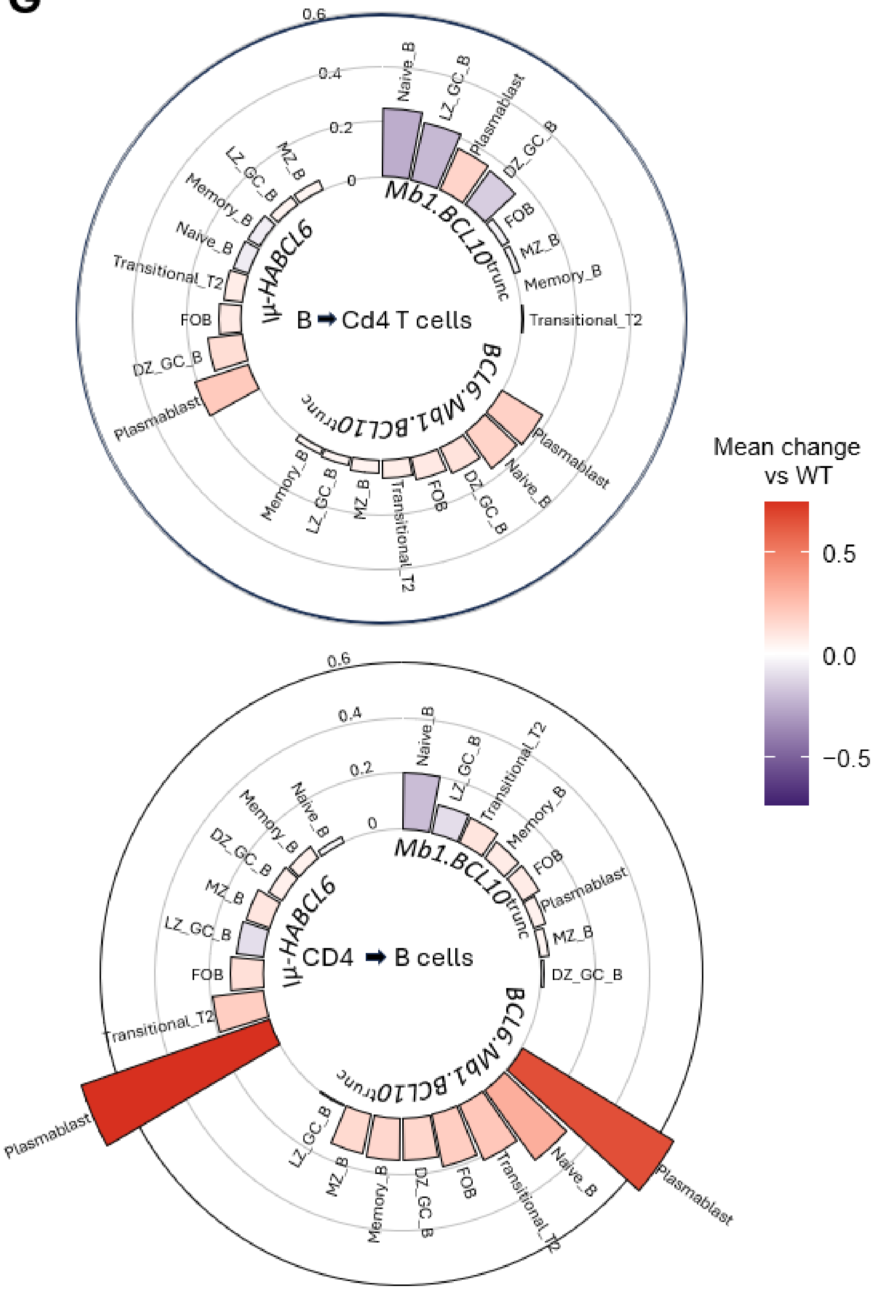
B ➡ Cd4 T cells
Mb1.BCL10trunc
BCL6.Mb1.BCL10trunc
Iμ-HABCL6
0
0.2
0.4
0.6
Naive_B
LZ_GC_B
Plasmablast
DZ_GC_B
FOB
MZ_B
Memory_B
Transitional_T2
CD4 ➡ B cells
Mean change
vs WT
0.5
0.0
−0.5

**H**

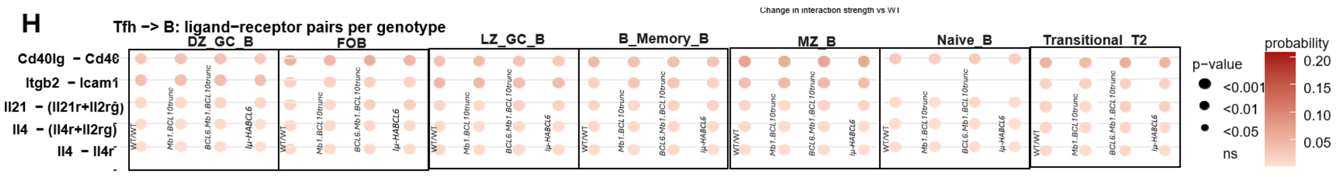
Tfh -> B: ligand−receptor pairs per genotype
Change in interaction strength vs WT
DZ_GC_B
FOB
LZ_GC_B
B_Memory_B
MZ_B
Naive_B
Transitional_T2
Cd40lg − Cd40
Itgb2 − Icam1
Il21 − (Il21r+Il2rg)
Il4 − (Il4r+Il2rg)
Il4 − Il4r
WT/WT
Mb1.BCL10trunc
BCL6.Mb1.BCL10trunc
Iμ-HABCL6
p-value
<0.001
<0.01
<0.05
ns
probability
0.20
0.15
0.10
0.05

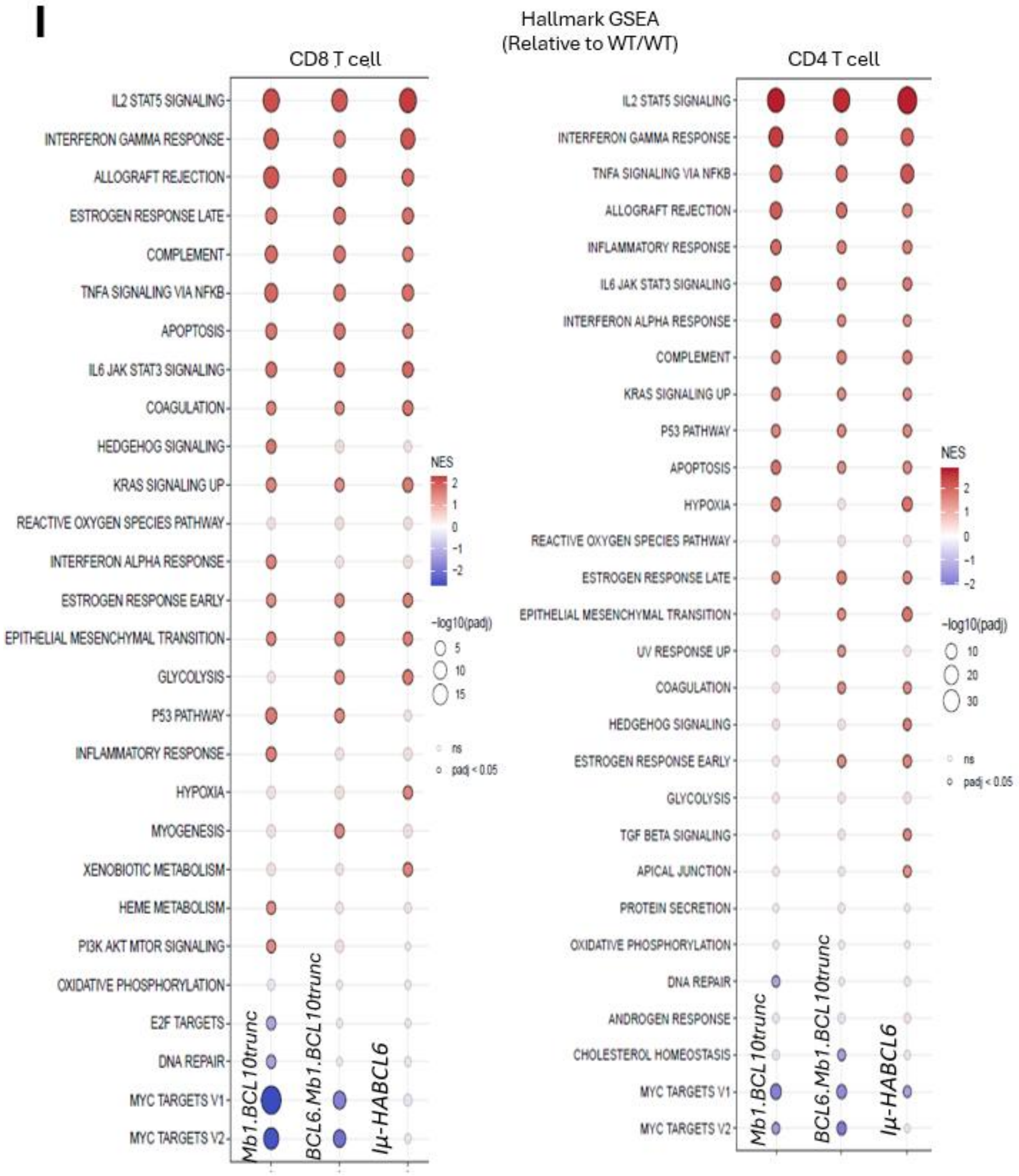
I
Hallmark GSEA
(Relative to WT/WT)
CD8 T cell
IL2 STAT5 SIGNALING
INTERFERON GAMMA RESPONSE
ALLOGRAFT REJECTION
ESTROGEN RESPONSE LATE
COMPLEMENT
TNFA SIGNALING VIA NFKB
APOPTOSIS
IL6 JAK STAT3 SIGNALING
COAGULATION
HEDGEHOG SIGNALING
KRAS SIGNALING UP
REACTIVE OXYGEN SPECIES PATHWAY
INTERFERON ALPHA RESPONSE
ESTROGEN RESPONSE EARLY
EPITHELIAL MESENCHYMAL TRANSITION
GLYCOLYSIS
P53 PATHWAY
INFLAMMATORY RESPONSE
HYPOXIA
MYOGENESIS
XENOBIOTIC METABOLISM
HEME METABOLISM
PI3K AKT MTOR SIGNALING
OXIDATIVE PHOSPHORYLATION
E2F TARGETS
DNA REPAIR
MYC TARGETS V1
MYC TARGETS V2
Mb1.BCL10trunc
BCL6.Mb1.BCL10trunc
Iμ-HABCL6
NES
2
1
0
-1
-2
-log10(padj)
5
10
15
ns
padj < 0.05
CD4 T cell
IL2 STAT5 SIGNALING
INTERFERON GAMMA RESPONSE
TNFA SIGNALING VIA NFKB
ALLOGRAFT REJECTION
INFLAMMATORY RESPONSE
IL6 JAK STAT3 SIGNALING
INTERFERON ALPHA RESPONSE
COMPLEMENT
KRAS SIGNALING UP
P53 PATHWAY
APOPTOSIS
HYPOXIA
REACTIVE OXYGEN SPECIES PATHWAY
ESTROGEN RESPONSE LATE
EPITHELIAL MESENCHYMAL TRANSITION
UV RESPONSE UP
COAGULATION
HEDGEHOG SIGNALING
ESTROGEN RESPONSE EARLY
GLYCOLYSIS
TGF BETA SIGNALING
APICAL JUNCTION
PROTEIN SECRETION
OXIDATIVE PHOSPHORYLATION
DNA REPAIR
ANDROGEN RESPONSE
CHOLESTEROL HOMEOSTASIS
MYC TARGETS V1
MYC TARGETS V2
Mb1.BCL10trunc
BCL6.Mb1.BCL10trunc
Iμ-HABCL6
NES
2
1
0
-1
-2
-log10(padj)
10
20
30
ns
padj < 0.05

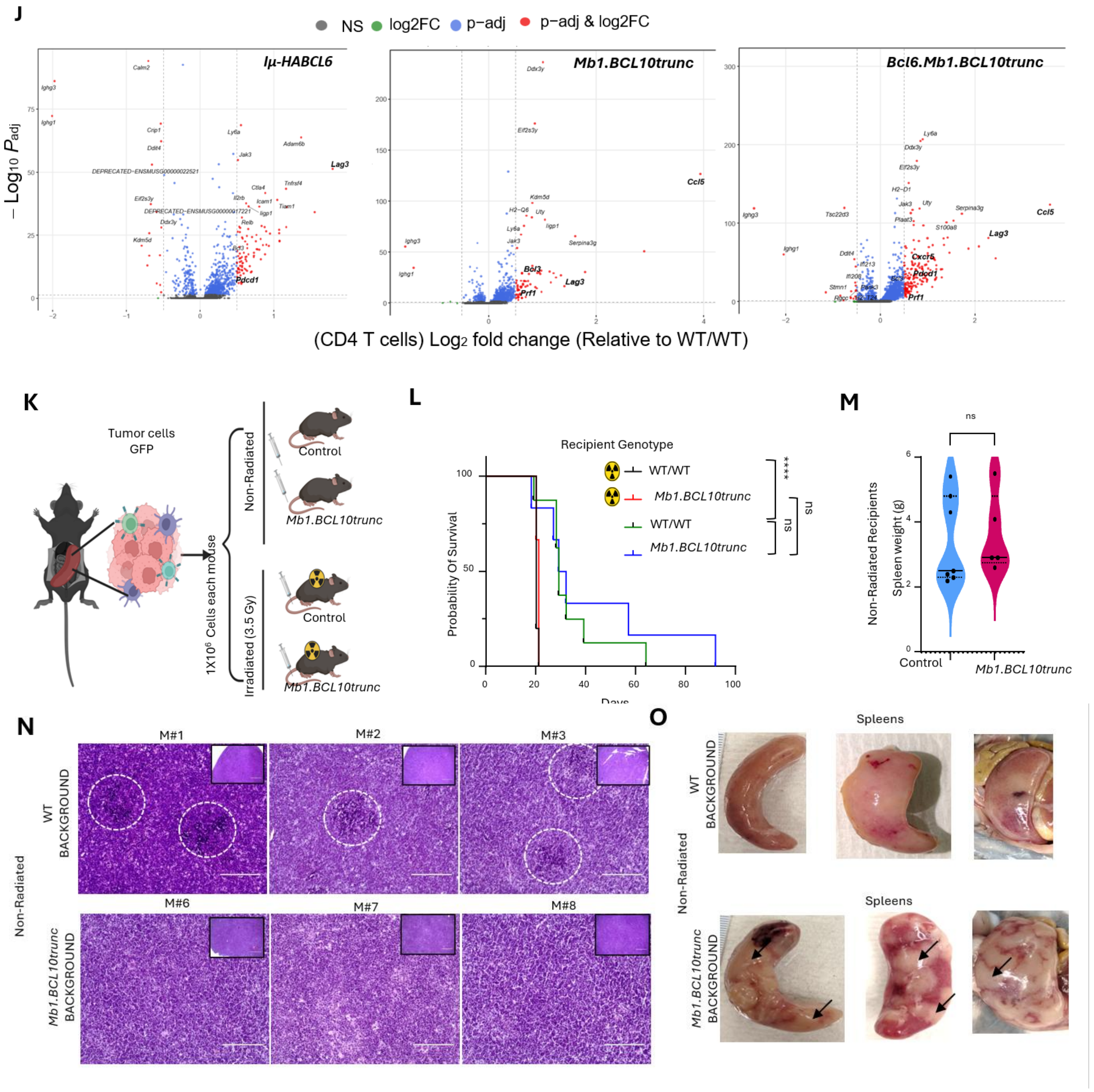


**Supplementary Figure 6. Single cell landscape of CD4 positive and CD8 positive population across genotypes; A.** UMAP projection of profiles from 26,950 CD4 positive T cells pooled form all genotypes (left). Cells are colored by annotated cluster: activated (CD4_activated), effector regulatory T cells (CD4_eTreg), central memory (CD4_Tcm), exhausted (CD4_Tex), T follicular helper (CD4_Tfh), Th1-like Tfh (CD4_Tfh_Th1), T follicular regulatory (CD4_Tfr), T helper1 (CD4_Th1), Interferon-stimulated gene high (CD4_Tisg), naïve (CD4_Tn), naïve with NFkβ (CD4_Tn_ NFkβ) and regulatory T cells (CD4_Treg). Smaller UMAPs show the same cells spits by genotype: WT/WT, *Mb1.BCL10*$^{trunc}$, *BCL6.Mb1.BCL10*$^{trunc}$, *Iμ-HABCL6* and *Cγ1.BCL10*$^{trunc}$. Red boxes highlight the CD4_Tex cluster, which is expanded in *Mb1.BCL10*$^{trunc}$ and

*BCL6.Mb1.BCL10*$^{trunc}$ relative to WT/WT, *Iμ-HABCL6* and *Cγ1.BCL10*$^{trunc}$. **B.** UMAP projection of profiles from 15,866 CD8 positive T cells pooled form all genotypes (left). Cells are colored by annotated cluster: central memory (CD8_Tcm), effector memory (CD8_Tem), Terminally differentiated effector memory T (CD8_Temra), exhausted (CD8_Tex), IFN-response T (CD8_Tisg), naïve (CD8_Tn), progenitor of exhausted T(CD8_Tpex). Smaller UMAPs show the same cells spits by genotype: WT/WT, *Mb1.BCL10*$^{trunc}$, *BCL6.Mb1.BCL10*$^{trunc}$, *Iμ-HABCL6* and *Cγ1.BCL10*$^{trunc}$. Red boxes highlight the CD8_Tex cluster, which is expanded in *Mb1.BCL10*$^{trunc}$ and *BCL6.Mb1.BCL10*$^{trunc}$ relative to WT/WT, *Iμ-HABCL6* and *Cγ1.BCL10*$^{trunc}$. **C.** Dot plots show expression of canonical marker genes across annotated CD8 positive (left) and CD4 positive (right) T cell clusters. Genes are grouped by Stacked bar plot showing proportions of CD4 positive and CD8 positive T cell population per genotype. **D.** Percentage of CD8 positive subsets across the genotypes. Bar plots show the percentage of the Tcm, Tem, Temra and Tpex across different genotypes. Genotypes were numbers as WT/WT (1), *Mb1.BCL10*$^{trunc}$ (2), *Cγ1.BCL10*$^{trunc}$ (3), *Iμ-HABCL6* (4) and *BCL6.Mb1.BCL10*$^{trunc}$ (5). Each dot represents one mouse. Bars indicate mean ±SEM (Wilcoxon test, $*P \leq 0.05$, $**P \leq 0.01$, $***P \leq 0.001$). **E.** Percentage of CD4 positive subsets across the genotypes. Bar plots show the percentage of the Tn, Tcm, Tn_ NFkβ, activated, Th1, Tfh, Tfh_Th1, Treg, eTreg, Tfr, Tex, Tisg across different genotypes. Genotypes were numbers as WT/WT (1), *Mb1.BCL10*$^{trunc}$ (2), *BCL6.Mb1.BCL10*$^{trunc}$(3), *Iμ-HABCL6* (4) and *Cγ1.BCL10*$^{trunc}$ (5). Each dot represents one mouse. Bars indicate mean ±SEM (Wilcoxon test, $*P \leq 0.05$, $**P \leq 0.01$, $***P \leq 0.001$). **F.** Lollipop plots show the change in interaction strength between B cell subsets and CD8 positive T cell subsets in *Mb1.BCL10*$^{trunc}$ (red), *BCL6.Mb1.BCL10*$^{trunc}$ (blue) and *Iμ-HABCL6* (green) mice relative to WT/WT mice. The left plot shows signaling from B (senders) cells to CD8 (receivers) positive T cells and the right plot shows signaling from CD8 (senders) positive T cells to B (receivers). **G.** Circular bar plots summarize the mean change in interaction strength between each B cell subset and CD4 positive T cells in *Mb1.BCL10*$^{trunc}$, *BCL6.Mb1.BCL10*$^{trunc}$ and *Iμ-HABCL6* mice relative to WT/WT mice. The top plot shows signaling from B (senders) cells to CD4 (receivers) positive T cells and the bottom plot shows signaling from CD4 (senders) positive T cells to B (receivers). Bar color and circle height indicates the mean change in strength of interaction. Red indicates increased interaction and purple indicates decreased interaction strength relative to WT/WT mice. **H.** Bubble plot shows selected ligand receptor pairs from CD4 positive Tfh cells (senders) to B cell subsets (receivers), predicted by CellChat across the different genotypes. Dot color indicates communication probability, and size indicates the *p*-value, determined by permutation test. **I.** Gene set enrichment analysis (GSEA) of hallmark gene sets in all CD8 positive (left) and all CD4 positive (right) from *Mb1.BCL10*$^{trunc}$, *BCL6.Mb1.BCL10*$^{trunc}$ and *Iμ-HABCL6* mice compared to WT/WT mice. Genes were ranked by log2fold from pseudobulk differential expression analysis. Dot color indicates normalized enrichment score, red indicates enrichment in mutant genotypes and blue indicates enrichment in WT. Dot size indicates -log10 (adjusted p value). The faded dots indicate non-significant values ($p$ adjusted $\geq 0.5$). **J.** Volcano plots show differentially expressed genes in CD4 positive T cells from *Iμ-HABCL6* (left), *Mb1.BCL10*$^{trunc}$ (middle), and *BCL6.Mb1.BCL10*$^{trunc}$ (right) compared with WT/WT mice, determined by pseudobulk/Wilcoxon test. The x-axis shows log2fold change, and the y-axis shows -log10 (adjusted *p* value). Dashed lines mark the significance thresholds, log2 FC>0.5 and p-adj<0.05. Dot colors: Gray shows the non-significant, Green shows the genes pass the fold change threshold only, blue indicates the genes pass the p-adj threshold only and red indicates the genes which passes both p-adj and fold change. Genes associated with exhaustion and cytotoxicity are labeled in bold. **K.** Experimental design for tumor transplantation experiment.

GFP positive tumor cells (BCL10 truncated protein) are isolated from the spleen of engrafted mice that were transplanted at 1 million cells per mouse into WT/WT background or *Mb1.BCL10*$^{trunc}$ background recipients, each either non-irradiated or sub-lethally irradiated (3.5 Gy) prior to transfer (n=5 mice per group). **L.** Survival of tumor bearing recipients. Kaplan-Meier curves show probability of survival of irradiated and non-irradiated WT/WT and *Mb1.BCL10*$^{trunc}$ mice following transplantation of tumor cells. Statistical significance was determined by the log-rank (Mantel-Cox) test (****$P \leq 0.0001$; ns, not significant). **M.** Spleen weights (g) of non-irradiated mice at the endpoints post-transplant. **N.** Representative H&E-stained spleen sections from non-irradiated WT/WT (top) and *Mb1.BCL10*$^{trunc}$ (bottom) recipients at end point. Dashed white circles outline the necrotic regions, which are present in WT/WT but largely absent in *Mb1.BCL10*$^{trunc}$ recipients. Insets show the lower-magnification overview of the whole section. Scale bars, 20 µm (large image) and 2000 µm (insets). **O.** Gross appearance of spleens from non-irradiated tumor recipients. Representative whole spleens from non-irradiated WT/WT (top) and *Mb1.BCL10*$^{trunc}$ (bottom) recipients at endpoint. Arrows indicate the pattern of defined tumor lesions, which are defined in *Mb1.BCL10*$^{trunc}$ but are largely diffused in WT/WT.

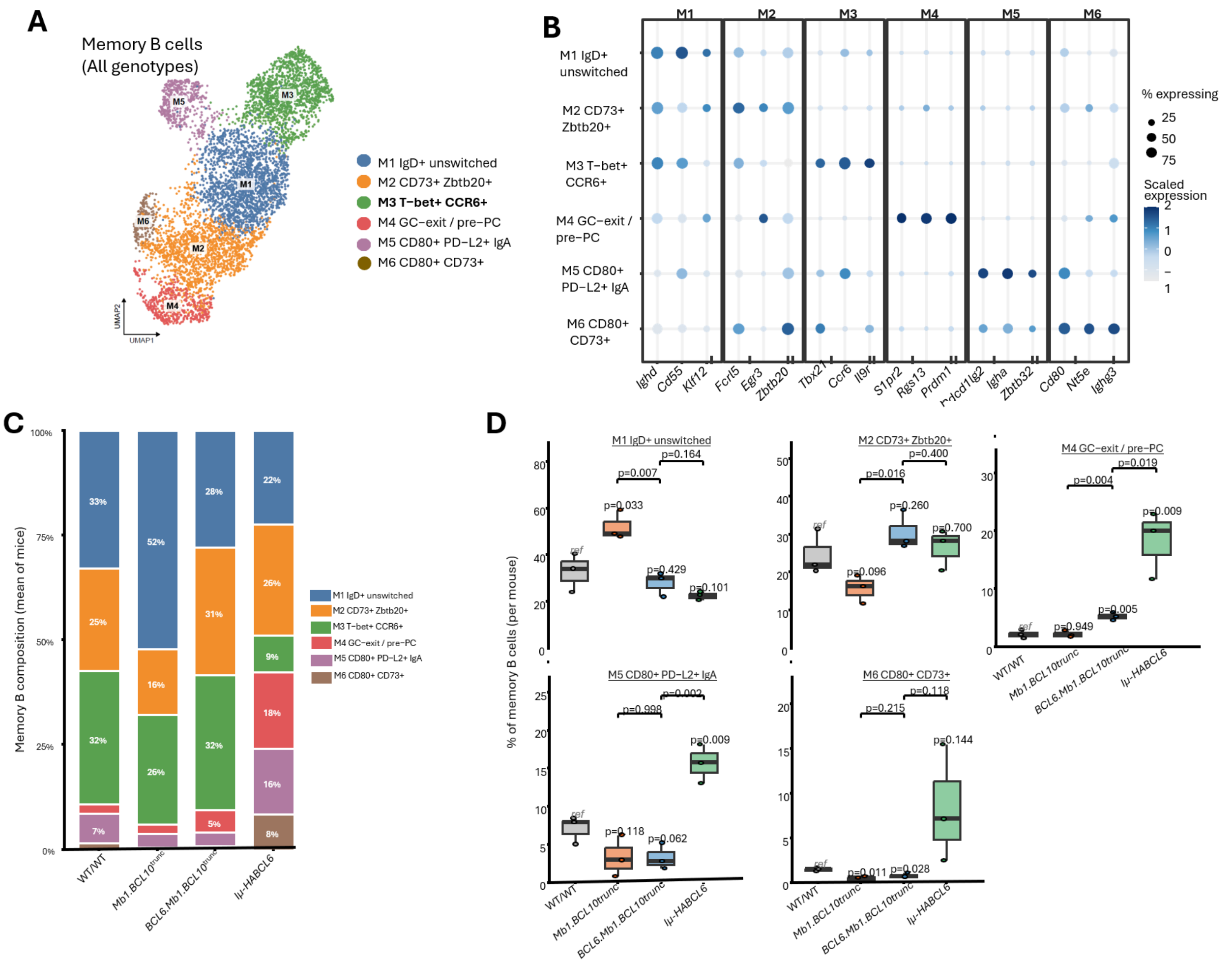


**Supplementary Figure 7. Transcriptional heterogeneity and genotype-dependent composition of splenic memory B cell in SRBC immunized mice; A.** UMAP projection of scRNA-seq profiles of memory B cells pooled from all genotypes colored by annotated subset:

M1($IgD^{+}$ unswitched), M2 ($CD73^{+}Zbtb20^{+}$ ), M3 ($CD80^{+}CD73^{+}$),M4 (GC exit/pre-plasma cell), M5 ($CD80^{+}PD\text{-}L2^{+}IgA^{+}$), and M6 ($CD80^{+}CD73^{+}$). (N=6251 cells; n=3 mice per genotype). **B**. Dot plot showing expression of subset-defining marker genes across the six memory B cells clusters. Dot size indicates the percentage of cells in each cluster expressing the gene, and color indicates scaled mean expression (z-score across clusters). **C**. Stacked bar plot showing the mean proportional composition of memory B cell subsets M1-M6 in WT/WT, *Mb1.BCL10*$^{\text{trunc}}$, *BCL6.Mb1.BCL10*$^{\text{trunc}}$ and *Iμ-HABCL6* mice . Bars represent the mean across mice within each genotype, with each subset's percentage indicated. **D**. Box plot showing the percentage of each memory B cell subset among total memory B cells, calculated per mouse, for subsets M1, M2, M4, M5 and M6. The M3 subset is shown in Fig 6. Boxes indicate the median and dot represents the mouse. WT/WT serves as reference group (ref).

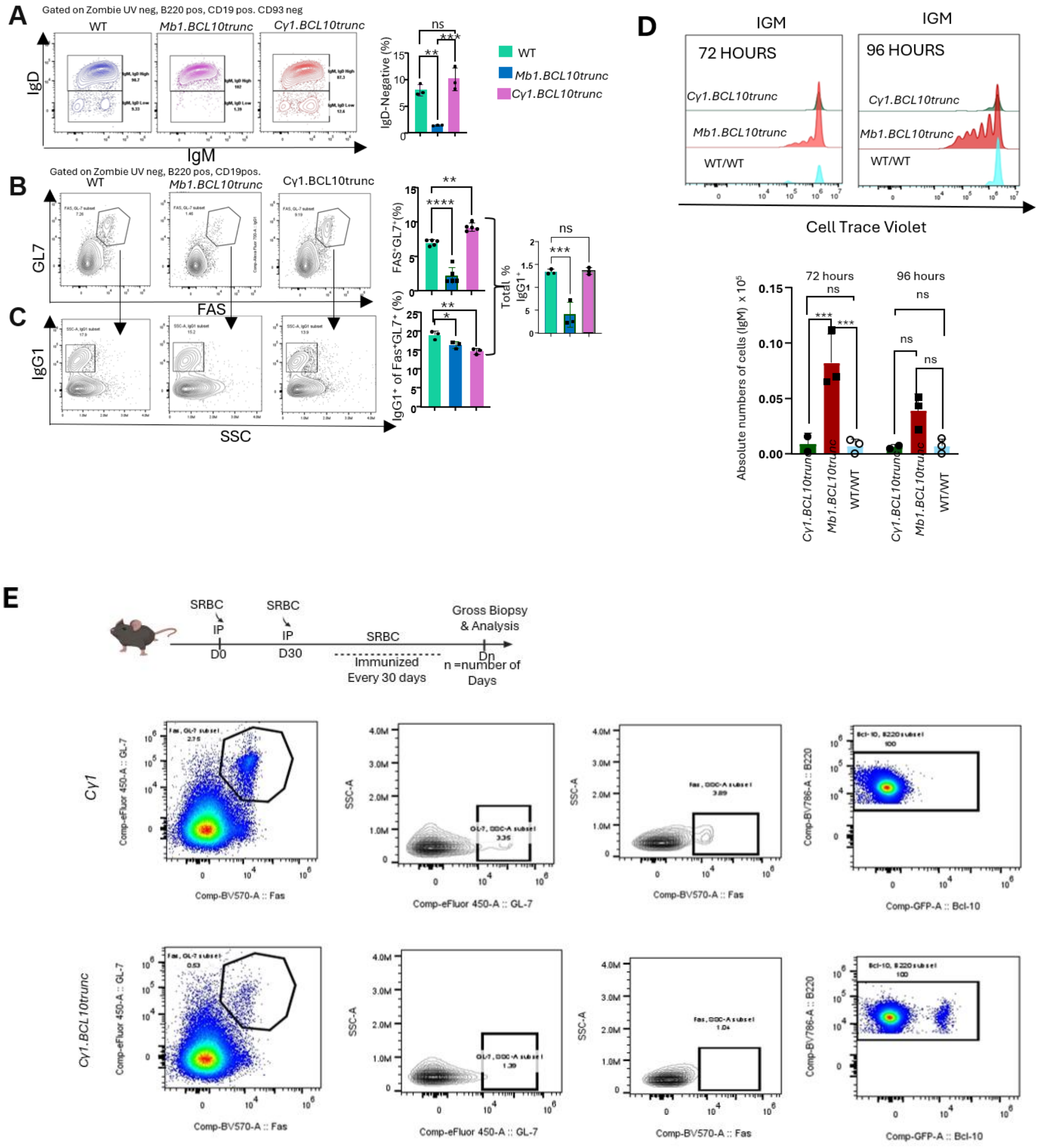

**Supplementary Figure 8. Repetitive SRBC immunization of *Cy1.BCL10*$^{\mathbf{trunc}}$ lead to abrogated GCs; A**. Representative flow plots of IgD versus IgM expression on live, mature B cells from WT/WT, *Mb1.BCL10*$^{trunc}$ and *Cy1.BCL10*$^{trunc}$ isolated 7 days post SRBC immunization. Gates define IgM,IgD$^{high}$ and IgM,IgD$^{low}$ populations, with the frequency of each shown. The bar graph quantifies the frequency of IgD negative cells among mature B cells. **B**. Representative contour plots of GL7 versus FAS on splenic B cells (gated on viable, B220, CD19 positive) from each genotype, with the FAS$^+$GL7$^+$ GC gate indicated and its frequency shown. The bar graph quantifies the frequency of double positive (FAS$^+$GL7$^+$) GC B cells. **C**. Representative plots of IgG1$^+$ versus Side scatter (SSC-A) on FAS$^+$GL7$^+$ GC B cells, IgG1$^+$ gate indicated. Bar graph quantifies the frequency of IgG1$^+$ cells among FAS$^+$GL7$^+$ GC B cells (left) and the frequency of total IgG1$^+$ cells among live, mature B cells. **D**. Representative histograms of CTV dilution in splenic mature B cells from WT/WT, *Mb1.BCL10*$^{trunc}$ and *Cy1.BCL10*$^{trunc}$ mice after 72 hours (left) and 96 hours (right) of αIgM (1µg/ml) stimulation ex-vivo condition. Successive peaks of reduced CTV intensity correspond to successive cell divisions. **E**. The representative FACS plot showing the FAS$^+$GL7$^+$ GC B cells (gated on viable, B220, CD19 positive) in serially immunized (SRBC, intraperitoneally) *Cy1.BCL10*$^{trunc}$ and control mice, *Cy1* . GFP corresponds to the expression of truncated BCL10 in *Cy1.BCL10*$^{trunc}$.The bar graph shows the absolute number of live B cells recovered after stimulation at each timepoint. For all bar graphs, each dot represents one mouse. Bars indicate mean ± SEM. Statistical significance was determined by one-way ANOVA with multiple comparison test;*p<0.05, **p<0.01, ***p<0.001,***p<0.0001 ;ns, not significant.
